# The SMYD3 methyltransferase promotes myogenesis by activating the myogenin regulatory network

**DOI:** 10.1101/804633

**Authors:** Roberta Codato, Martine Perichon, Arnaud Divol, Ella Fung, Athanassia Sotiropoulos, Anne Bigot, Jonathan B. Weitzman, Souhila Medjkane

## Abstract

The coordinated expression of myogenic regulatory factors, including MyoD and myogenin, orchestrates the steps of skeletal muscle development, from myoblast proliferation and cell-cycle exit, to myoblast fusion and myotubes maturation. Yet, it remains unclear how key transcription factors and epigenetic enzymes cooperate to guide myogenic differentiation. Proteins of the SMYD (SET and MYND domain-containing) methyltransferase family participate in cardiac and skeletal myogenesis during development in zebrafish, *Drosophila* and mice. Here, we show that the mammalian SMYD3 methyltransferase coordinates skeletal muscle differentiation *in vitro*. Overexpression of SMYD3 in myoblasts promoted muscle differentiation and myoblasts fusion. Conversely, silencing of endogenous SMYD3 or its pharmacological inhibition impaired muscle differentiation. Genome-wide transcriptomic analysis of murine myoblasts, with silenced or overexpressed SMYD3, revealed that SMYD3 impacts skeletal muscle differentiation by targeting the key muscle regulatory factor myogenin. The role of SMYD3 in the regulation of skeletal muscle differentiation and myotube formation, partially via the myogenin transcriptional network, highlights the importance of methyltransferases in mammalian myogenesis.

## INTRODUCTION

Myogenesis is a complex, stepwise process that is initiated by the commitment of precursor cells to the myoblast lineage, followed by cell-cycle arrest and gradual increase in muscle-specific gene expression, leading to terminal differentiation. The process culminates in the fusion of differentiated cells to generate myotubes, essential multinucleated syncytia that underlie the sarcomeric structure and contractile muscle functions. Muscle assembly and function rely on the coordinated expression of muscle-specific proteins that contribute both to muscle structure and to the control of excitation– contraction coupling, needed for force generation^1,2^. The combined and temporally-controlled action of the Myogenic Regulatory Factor (MRF) family of transcription factors (namely, MyoD, Myf5, Mrf4 and myogenin) together with selected epigenetic modifier enzymes generate highly-specific gene expression profiles required for myogenesis^3–5^. The MRF basic helix-loop-helix transcription factors bind to E-Box consensus sequences within the regulatory regions of muscle-specific genes^1,6^. While MyoD, Myf5 and Mrf4 act at the early onset of myogenesis to drive myoblasts formation, myogenin acts later in the cascade of myogenic activation, and is critical for the terminal differentiation of committed muscle cells and myofiber maturation^7–9^. Myogenin expression is not detected in proliferating myoblasts in culture, but appears early upon induction of differentiation. In *Myog*^*-/-*^ knockout mouse embryos, myoblasts are specified to muscle fate, but mice die at birth with severe skeletal muscle defects^10,11^. Similarly, loss of *myog* in zebrafish prevents myocytes fusion and formation of muscle fibers^12^. Genome-wide analysis has defined gene expression changes during myogenesis and the profiles of MRF binding upon differentiation ^3,13–15^. MyoD and myogenin regulate distinct, but overlapping, target genes and act sequentially at individual promoters^16,17^. Notably, MyoD alone is sufficient to fully activate the expression of early target genes (0-24h post-differentiation), whereas late-expressed genes (24-48h post-differentiation) require MyoD to initiate chromatin remodeling that subsequently facilitates myogenin binding and myogenin-mediated transcriptional activation^17^. MyoD can initiate the specification of muscle cell fate due to its capacity to recognize target genes within a native silent chromatin context and to initiate chromatin remodeling at these sites, allowing transcriptional activation^18–20^. Importantly, MyoD recruits most of the factors required to activate the *Myog* promoter upon differentiation, including histone methyltransferases (such as Set7/9), chromatin remodelers (like the SWI/SNF complex), as well as the basal transcriptional machinery via direct interaction with TAF3^20–22^.

Chromatin regulators drive major cell fate decisions, and histone lysine methyltransferases (KMTs) have emerged as key players in development, included cardiac and skeletal muscle formation ^23–25^. Aberrant regulation of these methylation events and alterations in global levels of histone methylation contribute to tumorigenesis and developmental defects^23^. However, our understanding of the role of epigenetic enzymes in myogenesis has lagged behind the characterization of the mechanistic contributions of the MRF transcription factors. The family of SMYD methyltransferases (SET and MYND domain-containing proteins) gained attention as novel myogenic modulators during development ^26,27^. For example, SMYD1, SMYD2 and SMYD4 play roles in cardiac and skeletal muscle differentiation in mouse, zebrafish and *Drosophila*, but the precise molecular mechanisms and relevant target genes are unknown^28–31^. The SMYD3 protein is frequently overexpressed in human cancers; it recognizes specific DNA sequences and has been reported to act as an epigenetic regulator through trimethylation of histone residues H3K4, H4K5 and H4K20 ^32–34^. SMYD3 is required for correct cardiac and skeletal muscle development in zebrafish ^35^. Proserpio and colleagues suggested a role for SMYD3 in regulating skeletal muscle atrophy via expression of *myostatin*, a strong negative regulator of muscle growth^36^. We studied the mechanisms of SMYD3 functions in myogenesis using *in vitro* myoblast differentiation. We investigated SMYD3 gain- and loss-of-function phenotypes and found that SMYD3 is required for the activation of the key MRF myogenin. Inhibition of SMYD3 expression or activity caused defective skeletal muscle differentiation and myotube formation, whereas SMYD3 overexpression enhanced differentiation and fusion. Transcriptome RNA-Seq analysis of mouse myoblasts upon SMYD3 knockdown (SMYD3_KD_) or SMYD3 overexpression (SMYD3_OE_) revealed a transcriptional network of genes involved in skeletal muscle structure and function. We show that SMYD3 acts upstream of a myogenin transcriptional program that is required for skeletal muscle differentiation.

## RESULTS

### SMYD3 overexpression enhances myogenic differentiation

Initial analysis showed that SMYD3 transcript and protein are expressed in proliferating, undifferentiated myoblasts and stably maintained throughout differentiation of either murine or human myoblasts (Supplementary information, Fig. S1A-D). To explore a role in myogenic differentiation, we overexpressed SMYD3 in C2C12 murine myoblasts using retroviral infections of HA-FLAG-tagged SMYD3. We generated two independent clonal cell lines, referred to as SMYD3 CL3 and SMYD3 CL5, and studied differentiation and myotube formation upon transfer to conventional differentiation media (DM). SMYD3-overexpressing (SMYD3_OE_) clones formed morphologically larger, multinucleated myotubes, compared to control cells (Fig. 1A-B). SMYD3 overexpression caused premature and elevated expression of differentiation markers, such as Muscle Creatine Kinase (MCK) and Myosin Heavy Chain (MyHC) compared to the controls (Fig.1C). RNA expression analysis revealed a marked upregulation of *Mck* and the fusion gene *Mymk*, as well as genes encoding components of the contractile apparatus (e.g. *Tnnc1, Myh3, Myl4, Atp2a1*) (Fig.1D and Supplementary information, S1E). Hence, SMYD3 overexpression enhanced myogenic differentiation and myotube formation, with premature appearance of muscle differentiation markers and the formation of larger, multinucleated myotubes. These results suggest that SMYD3 is a positive regulator of myoblast differentiation.

**Figure 1.**
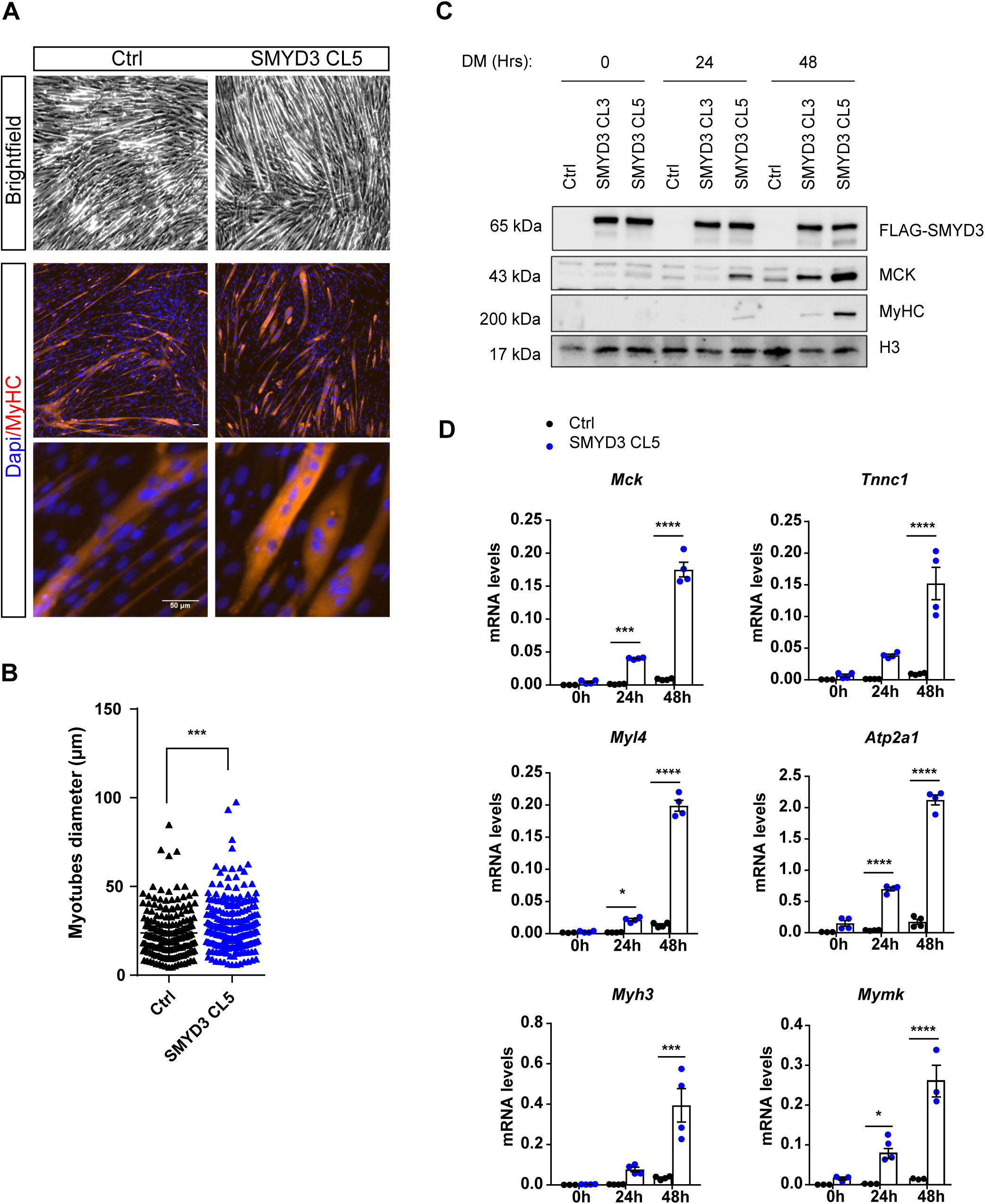
Smyd3 overexpression enhances C2C12 myoblast differentiation. **A)** Brightfield images and MyHC immunofluorescence analysis performed in differentiated C2C12 stable SMYD3_OE_ clones (SMYD3 CL5) and control empty vector (Ctrl). Cells were DAPI stained prior to immunofluorescence analysis. Scale bar 50µm. **B)** Quantification of myotubes diameter of >100 myotubes in four independent experiments, calculated within MyHC-expressing myotubes. Data are presented as average ± SEM. Mann-Whitney, *** p < 0.001. **C)** Time course analysis of protein expression during terminal differentiation of SMYD3_OE_ clones (SMYD3 CL3 and SMYD3 CL5) and control cells. Myoblasts were cultured in growth medium (GM) until they reached confluence, and then shifted to differentiation medium (DM). Cellular extracts were analyzed by western blot at 0h, 24h and 48h post-differentiation with antibodies against FLAG, muscle creatine kinase (MCK), and myosin heavy chain (MyHC). H3 is a loading control. Original blot images are shown in Supplementary Fig.S7A. **D)** qPCR analysis on Ctrl and SMYD3 CL5 SMYD3_OE_ cells at 0h, 24h and 48h post-differentiation in DM. mRNA levels of specific differentiation markers (*Mck, Tnnc1, Myl4, Myh3, Atp2a1, Mymk*) were normalized to *Gapdh* and *Rplp0* Ct values at the indicated timepoints. Graphs present means ± SEM of at least three independent experiments. ANOVA, * p < 0.05, ** p < 0.01, *** p< 0.001, **** p< 0.0001 vs. control respectively.

### SMYD3 knockdown impairs myogenic differentiation

To explore whether SMYD3 is required for skeletal myogenesis, we knocked-down SMYD3 expression in undifferentiated myoblasts by small interfering RNAs (siRNAs), and analyzed myogenic phenotypes. Knockdown of SMYD3 (SMYD3_KD_) severely impaired C2C12 differentiation; siSMYD3-transfected myoblasts remained predominantly as individual mononucleated cells, compared to the morphologically distinctive multinucleated myotubes in siControl (Fig. 2A). SMYD3 knockdown impaired myotube formation (even after 72h in DM), reduced the size and number of MyHC-positive cells, and decreased the fusion index and myotube diameter compared to control cells (Fig. 2A and B). SMYD3_KD_ cells exhibited significantly reduced levels of both MyHC and MCK proteins during a 3-day differentiation experiment (Fig.2C). Because the transcriptional landscape dramatically changes during the first 24 hours of myoblast differentiation^7^, we assessed whether SMYD3 silencing could impair transcription of the early myogenic cascade. We analyzed mRNA expression of myogenic differentiation factors in early differentiating C2C12 cells upon SMYD3 silencing. siSMYD3 significantly attenuated the transcriptional activation of myogenic markers (e.g. *Mck, Tnnc1, Myh3, Myl4, Atp2a1*, and *Mymk*) at 24h post-differentiation, compared to control cells (Fig.2D). These observations were confirmed with a second independent siRNA oligo, si*SMYD3*#2 (Supplementary information, Fig. S2A-E). We observed a similar delay in myogenic differentiation in primary murine myoblasts, with decreased MCK and MyHC protein levels in SMYD3_KD_ cells, compared to siCtrl controls (Supplementary information, Fig. S2F). These data demonstrate that transient SMYD3 knockdown prevented correct myogenic differentiation in murine myoblasts, delayed expression of key muscle differentiation markers and strongly impaired myoblast fusion. Additional experiments with human immortalized myoblasts, showed that SMYD3 is expressed in both undifferentiated and differentiated human cells (Supplementary information, Fig. S1C and D). siRNA knockdown of human *SMYD3* also impaired myogenic differentiation, reducing protein levels of MCK and MyHC markers (Supplementary information, Fig. S3A). We extended these results by stable and sustained *SMYD3* silencing using CRISPRi technology^37^. Again, *SMYD3*-silencing (sgSMYD3) caused a delay in myogenic differentiation, and marked downregulation of MCK protein levels compared to controls (sgCtrl) (Supplementary information, Fig. S3B-C) with a significant reduction in *MCK* and muscle actin *ACTA1* RNA levels (Supplementary information, Fig. S3D). To rule out clonal SMYD3_OE_ effects, we performed additional experiments to show that siSMYD3 reversed MyHC staining, differentiation marker expression, myotube size and the differentiation potential of SMYD3_OE_ cells (Supplementary information, Fig. S3E-G). These results exclude possible clonal effects, and showed that SMYD3 knockdown could counteract the enhancement of myogenesis observed in SMYD3_OE_ clones. Other members of the SMYD protein family of lysine methyltransferases including SMYD1, 2 and 4 are implicated in skeletal and cardiac differentiation in different model organisms ^28,30,31^. We tested whether *Smyd1, Smyd2* and *Smyd4* mRNA levels were affected in SMYD3_KD_ cells. Interestingly, we found that only the expression of the multifunctional myogenic activator *Smyd1* was significantly reduced upon SMYD3 silencing (Supplementary information, Fig. S3H). Likewise, *Smyd1* transcript levels were also strongly upregulated in SMYD3_OE_ cells, whereas mRNA levels of both *Smyd2* and *Smyd4* were unaffected (Supplementary information, Fig. S3H). Overall, these loss-of-function experiments demonstrate that SMYD3 is required for appropriate myogenic transcriptional activation, myoblast fusion and myotube formation. SMYD3_KD_ in myoblasts caused deregulated expression of differentiation markers and defective terminal differentiation.

**Figure 2.**
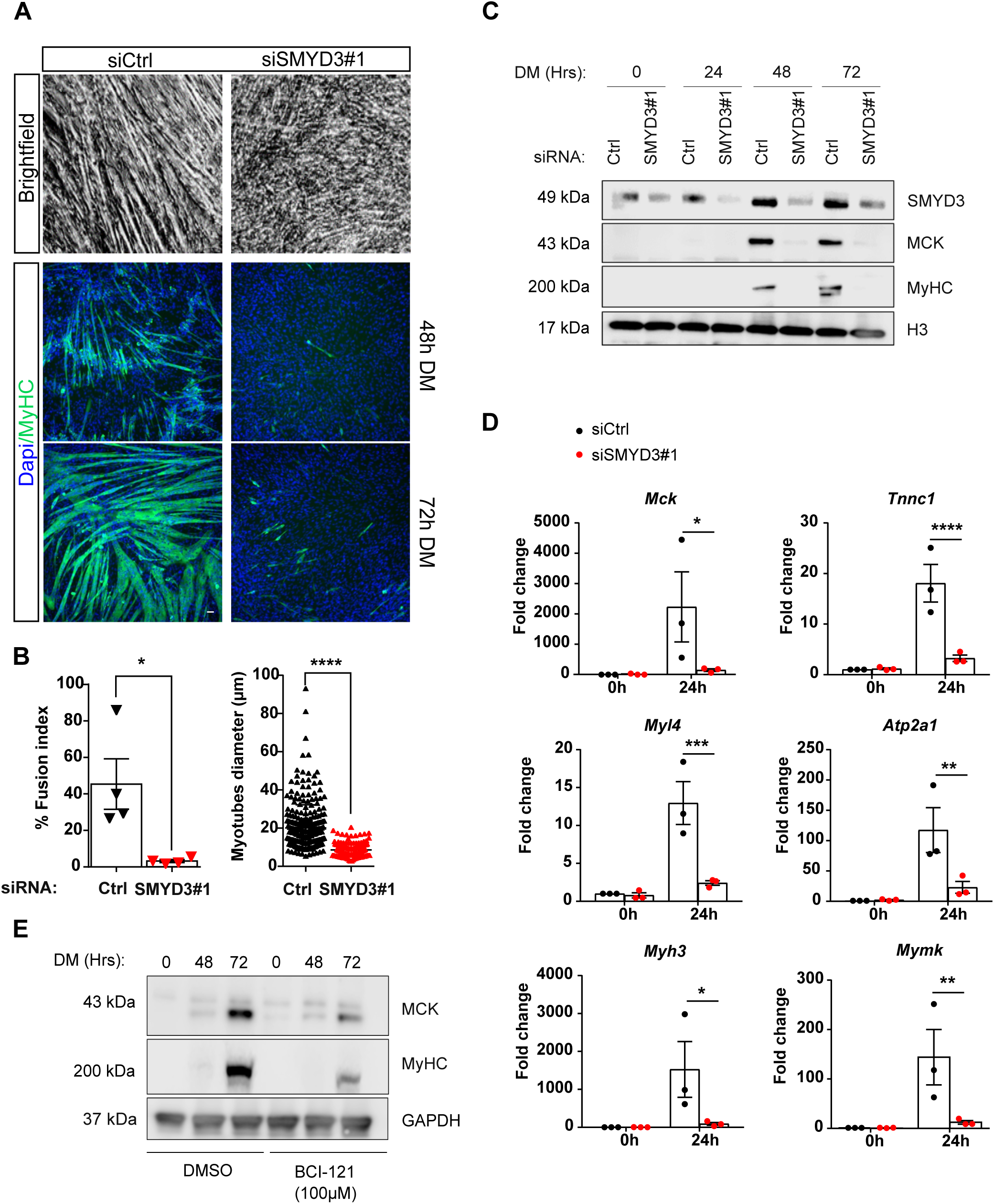
Knockdown of Smyd3 or its pharmacological inhibition impair skeletal muscle differentiation. **A)** Brightfield images and MyHC immunofluorescence analysis performed in differentiated in C2C12 myoblasts after transfection with control or *Smyd3* siRNAs. Cells were transferred to differentiation medium and stained with MyHC antibody at the indicated timepoints. Cells were Hoechst stained prior to immunofluorescence analysis. Scale bar, 50µm. **B)** Left panel: quantification of fusion index of four independent experiments calculated as percentage of nuclei within MyHC-expressing myotubes at 72h in presence of the various siRNA. Data shown as mean ± SEM. Mann-Whitney, * p < 0.05. Right panel: Quantification of myotubes diameter of >100 myotubes in four independent experiments, calculated within MyHC-expressing myotubes. Data are presented as average ± SEM. Mann-Whitney, **** p < 0.0001. **C)** Time course of protein expression during terminal differentiation of SMYD3_KD_ cells. Cellular extracts were analyzed by western blot at 0h, 24h, 48h and 72h in DM with antibodies against SMYD3, muscle creatine kinase (MCK), and myosin heavy chain (MyHC). H3 is a loading control. Original blot images are shown in Supplementary Fig.S7B. **D)** qPCR analysis on C2C12 cells after transfection with control or *Smyd3* siRNAs. mRNA levels of *Smyd3* and specific differentiation markers (*Mck, Tnnc1, Myl4, Myh3, Atp2a1, Mymk*) were normalized to *Gapdh* and *Rplp0* Ct values at the indicated timepoints. Data shown as mean fold change ± SEM of three independent experiments. ANOVA, * p < 0.05, ** p < 0.01 **** p<0.001 vs. control respectively. **E)** C2C12 cells were treated with control DMSO or 100µM of BCI-121 and differentiated for 72h. Cellular extracts were analyzed by western blot at the indicated time points with antibodies against muscle creatine kinase (MCK), and myosin heavy chain (MyHC). GAPDH is a loading control. Original blot images are shown in Supplementary Fig.S7F.

### Pharmacological inhibition of SMYD3 decreases myogenic differentiation potential

BCI-121 is a novel SMYD3 small-molecule inhibitor that specifically reduces SMYD3 activity, by competing with histones for binding to SMYD3 and preventing its recruitment to the promoter region of target genes^38^. To explore further SMYD3 function in myogenesis, we cultured and differentiated C2C12 cells with medium that contained 100µM of BCI-121, or DMSO controls, and analyzed MyHC and MCK markers expression by western blot analysis. BCI-121 treatment significantly decreased the differentiation efficiency, recapitulating the effects observed upon transient SMYD3 knockdown (Fig.2E). This inhibitory effect was also validated in SMYD3_OE_ cells (Supplementary information, Fig. S3I). Hence, SMYD3 pharmacological inhibition using the BCI-121 inhibitor significantly reduced myoblast differentiation potential, supporting a role for SMYD3-mediated methylation in myogenesis.

### SMYD3 does not influence myoblast proliferation

Myogenic terminal differentiation requires a delicate balance between proliferation and differentiation, involving permanent exit from the cell cycle and concomitant activation of a muscle-specific genetic program. Consequently, factors that regulate cell cycle progression can inhibit myogenic differentiation by preventing cell cycle exit ^39^. SMYD3 is frequently overexpressed in cancer cells and enhances cell proliferation ^32^. It was therefore important to determine whether SMYD3-dependent effects on muscle differentiation result from altered proliferative signals and impaired cell cycle arrest upon mitogen withdrawal. We observed no differences in the global number of cells in SMYD3_KD_ or SMYD3_OE_ myoblasts following induction of differentiation, compared to controls. To evaluate the impact of SMYD3 on myoblast proliferation and cell cycle arrest, we performed EdU incorporation analysis in C2C12 cells and measured the percentage of EdU-positive nuclei in growth medium (GM) and after 24h in Differentiating media (DM). These experiments showed comparable percentages of EdU-positive cells in both SMYD3_OE_ and SMYD3_KD_ cells and controls, in both proliferating and differentiating conditions (Fig.3A-B and Supplementary information, Fig. S4A-B). Furthermore, Cyclin D1 and cyclin-dependent kinase inhibitor 1 p21 protein levels were not affected by SMYD3_KD_ or SMYD3_OE_ (Fig.3C-D). Thus, SMYD3 is not implicated in cell cycle regulation in myoblasts and SMYD3-associated differentiation phenotypes are uncoupled from cell cycle exit.

**Figure 3.**
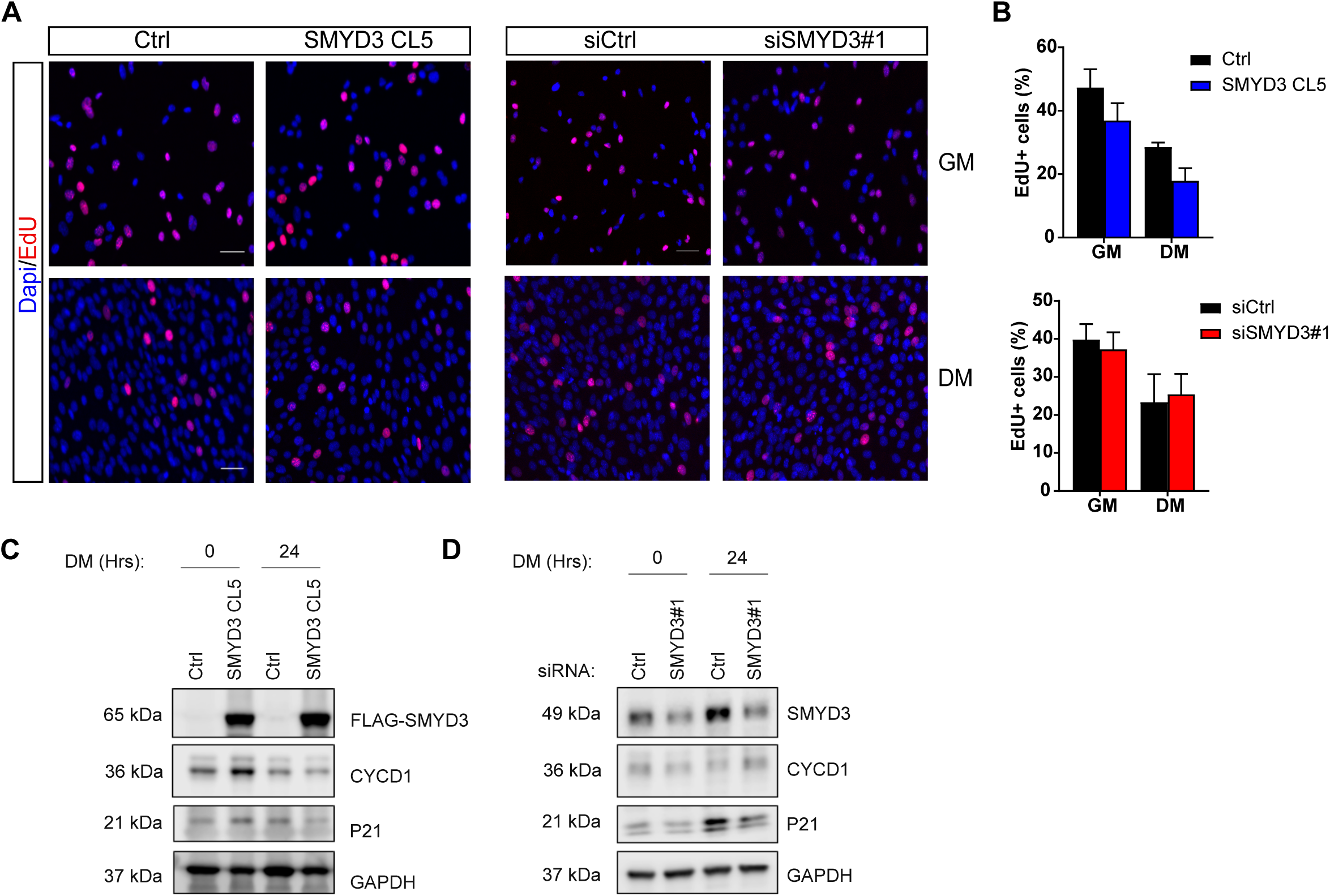
SMYD3 does not affect myoblasts proliferation and cell cycle. **A)** EdU staining was performed in SMYD3_OE_ cells (SMYD3 CL5) and control cells (Ctrl) (left panel) and in SMYD3_KD_ (SMYD3#1) or a scramble sequence (siCtrl) (right panel). In both cases, cells were labeled with EdU either in GM or after 24h in DM. Pictures are one representative of 3 independent experiments. EdU (red) and DAPI (blue) staining (scale bar: 50µm). **B)** Percentage of EdU positive nuclei/DAPI labeled nuclei in randomly selected fields, in SMYD3_OE or_ SMYD3_KD_ cells, compared with relative controls. Error bars represent SEM from quantifications of 3 independent experiments. **C)** Western blot analysis of SMYD3, CYCD1 and p21 protein expression during terminal differentiation of SMYD3_OE_ (SMYD3 CL5) and control cells (Ctrl). Original blot images are shown in Supplementary Fig.S7C. **D)** Analysis of SMYD3_KD_ (siSMYD3#1) or scramble sequence (siCtrl) cells. GAPDH is used as internal control. Original blot images are shown in Supplementary Fig.S7D.

### SMYD3 regulates genes associated with myogenic differentiation

SMYD3 has been linked to active histone marks associated with euchromatin formation and we observed that SMYD3 levels in myoblasts affect expression of key differentiation markers early upon muscle differentiation. To define SMYD3-dependent molecular pathways and target genes, we performed RNA-Seq analysis of undifferentiated (0h) and differentiated (24h) C2C12 myoblasts SMYD3_KD_ or SMYD3_OE_ cells. We compared two different siRNA SMYD3 (siSMYD3#1 and siSMYD3#2) with control siRNA (siCtrl). And we compared SMYD3_OE_ SMYD3 CL5 cells with control cell lines. We identified 5013 genes (1096 genes up and 3917 genes down) that were differentially expressed between undifferentiated (0h) and differentiated (24h) C2C12 siCtrl myoblasts. Gene Ontology (GO) analysis of differentially expressed genes confirmed enrichment in skeletal muscle development and cell cycle progression (not shown). Among the 5013 genes whose expression changes upon differentiation at 24h in the control myoblasts, we found 311 genes whose expression appeared to be SMYD3-dependent (107 upregulated and 204 downregulated genes) (Supplementary information, Table S1). We performed enrichment analyses using the Gene Ontology Enrichment Analysis tool (http://geneontology.org/). Downregulated genes were enriched for GO terms associated with skeletal muscle development and muscle contraction, whereas upregulated genes were enriched for cell adhesion, cell division and cell migration genes. Conversely, SMYD3_OE_ cells show 836 genes whose expression appeared to be SMYD3-dependent at 24h post differentiation (412 upregulated and 424 downregulated genes) (Supplementary information, Table S2). These upregulated genes highlighted skeletal muscle development, actin organization, sarcomeric structures and muscle contraction processes. Downregulated genes were mainly related to cell motility, cell adhesion and cell communication and signaling. To focus on potential SMYD3 target genes, we identified common deregulated genes between SMYD3_KD_ and SMYD3_OE_ cells at 24h post-differentiation. These 72 differentially-expressed genes exhibited a strong anti-correlation between SMYD3-loss and gain of function cell systems (Fig. 4A and Supplementary information, Table S3) and were enriched for genes involved in skeletal muscle development, contraction and cell migration or positioning (Fig. 4B). Further analysis revealed three main classes of biological functions: (i) gene families coding for structural sarcomeric proteins required for myofibril assembly, for example myosin light chain (*MYL1, MYL4, MYL6B, MYLPF*), myomesins (*MYOM1, MYOM2,MYOM3*) and Troponins (*TNNI1, TNNI2, TNNT1, TNNT3*), (ii) genes encoding calcium pumps and calcium binding proteins required for muscle excitation and contraction (e.g. *ATP2A1, Sarcalumenin, RYR1, RYR3*), and (iii) genes encoding proteins involved in cell-extracellular matrix interactions (e.g. *MMP10, SGCA*). The list also includes three genes involved in myoblast fusion; the muscle-specific Ser/Arg-rich protein specific kinase 3 (*SRPK3*), the Kelch-like protein 41 (*KLHL41*) and the myoblast fusion protein *Myomaker (Mymk).* Interestingly, we observed deregulated expression of the MRF *myogenin*, a key regulator of skeletal muscle differentiation; *myogenin* was markedly reduced in SMYD3_KD_ cells and significantly elevated in SMYD3_OE_ cells, at 24h post-differentiation compared to control cells. Notably, myogenin was the only transcription factor on this list of differentially expressed genes. Hence, the genome-wide expression profiling of differentiating C2C12 cells suggested that SMYD3 could participate in an early cascade of transcriptional activation (0-24h) and that genes important for muscle formation and activity are impacted by SMYD3. Our analysis also raised the possibility that deregulated *myogenin* expression might account for the defects in muscle gene expression and differentiation. Myogenin target genes have been widely studied^3,17,40–42^. To test this hypothesis, we compared the SMYD3-dependent differentially expressed genes with a published list of myogenin target genes reported for myogenin knockdown studies (#GSE19967)^43^. We identified 894 genes differentially expressed upon myogenin knockdown in C2C12 cells at 24h post-differentiation (p-value cut-off of 0.01, following ID microarray conversion) (Supplementary information, Table S4). Of these, 37 genes were differentially regulated in our SMYD3_KD_ dataset (i.e. 11.8% of differential genes in siSMYD3 C2C12 at 24h post-differentiation), and this enrichment is statistically significant (p = 8.19 e-17 by hypergeometric distribution test). Conversely, 41 of these myogenin-dependent genes were in common with our SMYD3_OE_ C2C12 dataset (i.e. 12.3% of SMYD3_OE_ C2C12 genes) (Supplementary information, Table S4). Again, this enrichment is significant (p = 1.78 e-06 by hypergeometric distribution test). Thus, the expression of myogenin-regulated genes correlated with SMYD3_KD_, and were anti-correlated with SMYD3_OE_ datasets (Fig. 4C and Supplementary information, Fig. S5C-D). Surprisingly, only 11 genes were in common within the three datasets, including *myogenin* itself and the myogenin target *Mymk* (Fig. 4D-E). Our findings show that SMYD3-differentially expressed genes display a myogenin-transcriptional signature and support a model in which the deregulated transcriptional program in SMYD3_KD_ and SMYD3_OE_ cells is due to defective activation of the myogenin pathway.

**Figure 4.**
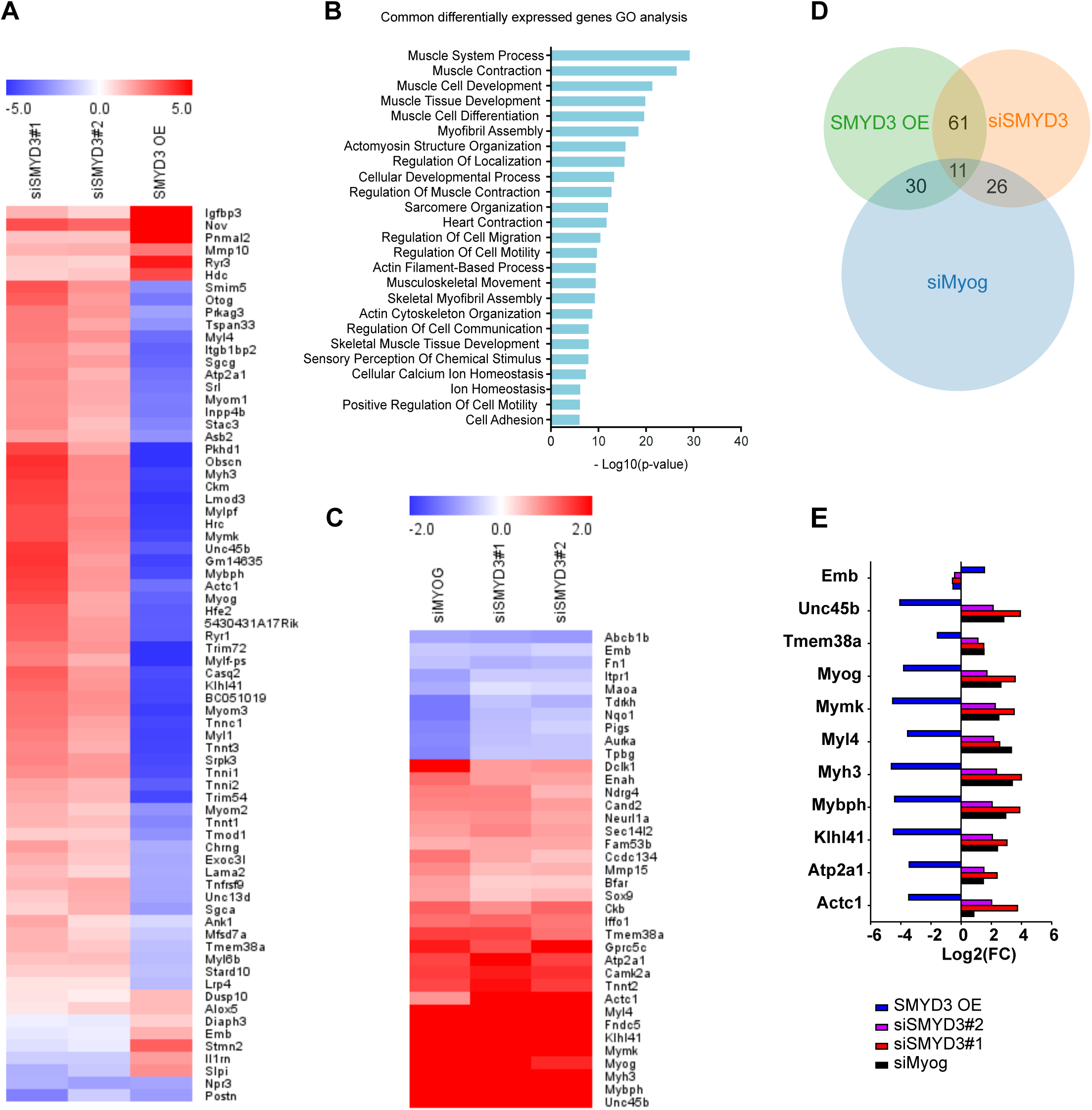
Transcriptome analysis of SMYD3_KD_ and SMYD3_OE_ myoblasts. **A)** Heat map of common DE genes (p-value <0.01) in SMYD3_KD_ and SMYD3_OE_ cells at 24h post-differentiation (Log2 FC, hierarchical clustering using Euclidean distance). **B)** PANTHER GO enrichment analysis tool was used to identify significant GO terms (Biological Process) enriched in the list of common DE genes between SMYD3_KD_ and SMYD3_OE_ cells at 24h post-differentiation. **C)** Heat map of common DE genes (p-value <0.01) in SMYD3_KD_ and siMyog C2C12 cells, from the study of Liu et al, 2010, at 24h post-differentiation (Log2 FC, hierarchical clustering using Euclidean distance). **D)** Venn diagram representing the number of common genes between SMYD3_KD_, SMYD3_OE_ and siMyog from the study of Liu et al, 2010, in C2C12 at 24h post differentiation. **E)** Histogram representing Log2(FC) levels of the 13 genes found in common between the present study and siMyog from the study of Liu et al, 2010.

### SMYD3 is necessary to induce *myogenin* expression upon differentiation

Our study showed SMYD3-dependent changes in *myogenin* expression levels following SMYD3 silencing or overexpression in murine and human myoblasts (Fig.5A-B, Supplementary information Fig.S6A). Furthermore, aberrant activation of Myogenin expression was also observed after SMYD3 pharmacological inhibition, both in wild-type C2C12 and in SMYD3 overexpressing cells (Supplementary information Fig.S6B). *Myogenin* expression is tightly regulated during differentiation through chromatin modifications on its promoter and a positive autoregulation loop^41^. Publicly available chromatin immunoprecipitation (ChIP) sequencing (ChIP-seq) ENCODE data showed that the *myogenin* locus is enriched for H3K4me3 upon myogenic activation and that myogenin binds to its own promoter upon differentiation (Fig. 5C). SMYD3 was reported to control gene expression through H3K4 di/tri-methylation of the promoters of specific target genes^32^. We performed ChIP analysis of H3K4me3 levels on the *myogenin* locus to assess whether SMYD3 affects histone methylation levels. SMYD3_KD_ cells exhibited decreased H3K4me3 levels on the *myogenin* gene upon differentiation, whereas promoter and distal regions were not affected (Fig.5D-E). Conversely, H3K4me3 levels increased significantly on the same region in SMYD3_OE_ cells compared with controls (Fig.5F). SMYD3 can bind to DNA containing 5’-CCCTCC-3’ or 5’-GAGGGG-3’ sequences on the regulatory regions of target genes^32^, and analysis of the *myogenin* locus revealed the presence of multiple SMYD3 binding sites, both upstream and downstream of the *myogenin* transcription start site (TSS). However, we could not detect any significant binding of SMYD3 to the *myogenin* locus (data not shown). Our data suggest that although SMYD3 is required for *myogenin* transcriptional activation and is associated with H3K4me3 changes, SMYD3 action might still be indirect. Activation of *myogenin* expression at early stages of skeletal muscle differentiation is essential for myogenic progression and myogenin controls the expression of late muscle genes by cooperating with other transcription factors^41^. Therefore, we tested whether exogenous *myogenin* expression could rescue the myogenic potential of SMYD3_KD_ C2C12. Myogenin ectopic expression partially rescued myotubes formation in siSMYD3 cells, (Fig.5G-H, Supplementary information Fig.S6C). Overall, these findings show that SMYD3 is required for myoblast differentiation through activation of *myogenin* transcriptional program and also via a myogenin-independent pathway.

**Figure 5.**
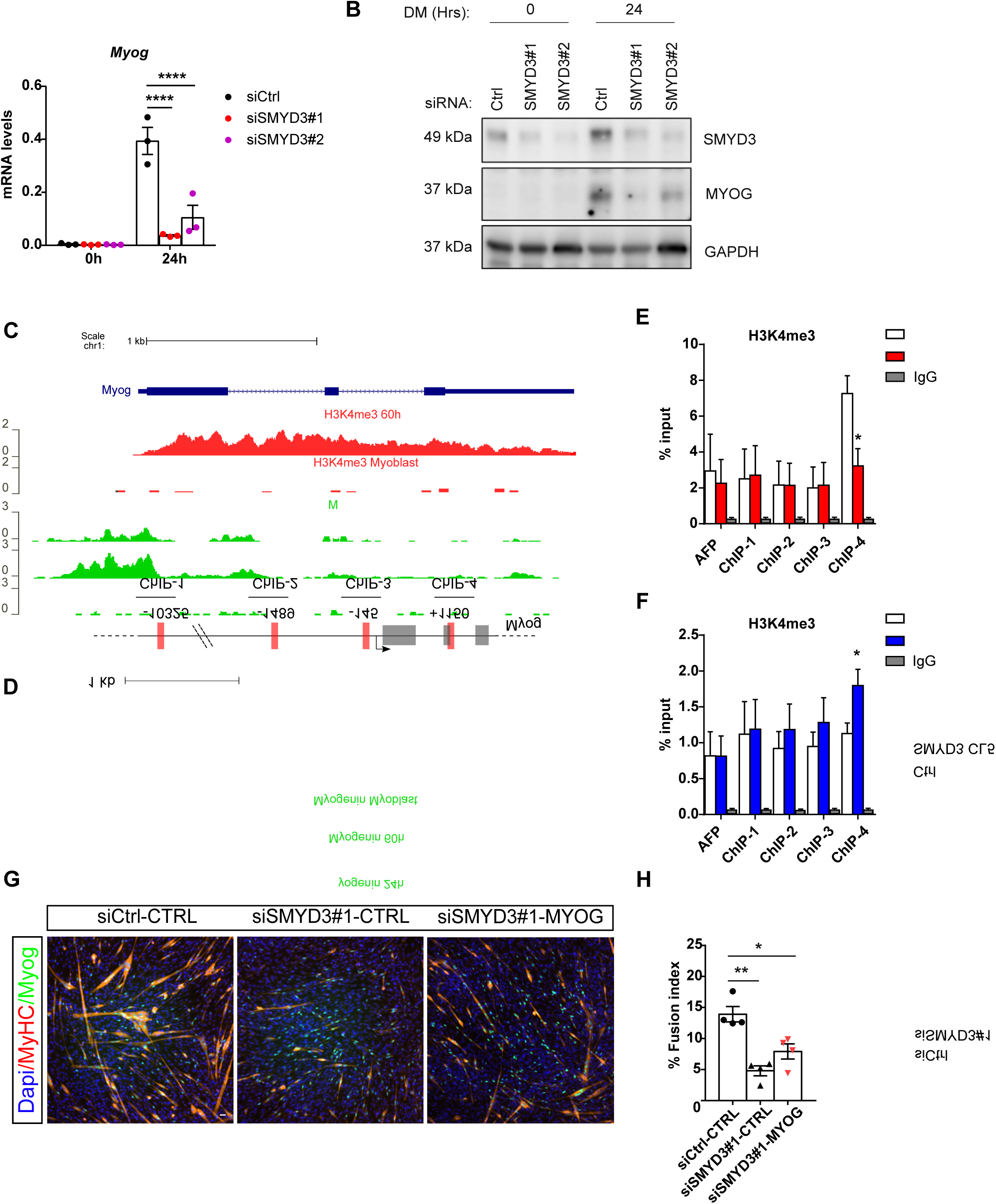
SMYD3 regulates *myogenin* expression. **A)** Relative expression levels of *myogenin* transcripts monitored by qRT-PCR in siCtrl and siSMYD3 (siSMYD3#1 and siSMYD3#2) undifferentiated (0h) and differentiated (24h) C2C12 cells. Data shown as means ± SEM of three independent experiments. ANOVA, **** p< 0.0001 vs. control respectively. **B)** Western blot showing decreased expression of myogenin in undifferentiated (0h) and differentiating (24h) C2C12 cells after *Smyd3* knockdown. GAPDH was used as an internal control. Original blot images are shown in Supplementary Fig.S7E. **C)** UCSC Genome Browser snapshot of H3K4me3 and myogenin ChIP-seq reads, aligned along *Myog* locus, in undifferentiated myoblasts and after 24h or 60h of differentiation in C2C12. **D)** Schematic representation of murine *Myog* locus, illustrating exons (gray rectangles), introns and intergenic regions (black line). Regions amplified by the primers used in the ChIP assays are represented as red rectangles, and their position on the locus is indicated. Scale bar (1 Kb) is indicated above. **E)** ChIP assay of chromatin from C2C12 at 24h of differentiation, after transient transfection with a control siRNA (siCtrl) or an siRNA targeting SMYD3 (siSMYD3#1). Immunoprecipitation was performed using H3K4me3 antibody or IgG, AFP was used as negative control position, ChIP-qPCR n=3. Unpaired two-tailed t-test, *p< 0.05 vs. control respectively. **F)** ChIP assay was performed with chromatin from SMYD3 CL5 myoblasts and control cell line (Ctrl), after 24h in DM. Immunoprecipitation was performed using H3K4me3 antibody or IgG, AFP was used as negative control position, ChIP-qPCR n=6. Unpaired two-tailed t-test, * p< 0.05 vs. control respectively. **G)** Immunofluorescence analysis of myogenin and MyHC in siRNA-treated C2C12 cells at 72h in DM, following overexpression of myogenin or a control vector. Scale bar 50µm. **H)** Fusion index was calculated by immunofluorescence staining as the percentage of nuclei within MyHC expressing myotubes, in siCtrl or siSMYD3 cells, after overexpression of myogenin or control empty vector. Data shown as means ± SEM of four independent experiments. ANOVA, ** p< 0.01, * p< 0.05.

## DISCUSSION

Studies of SMYD3 methyltransferase established a role in promoting cancer cell proliferation, invasion and metastasis. Some of these cancer phenotypes were attributed to regulation of specific target genes including *WNT10B*^44^, *hTERT*^45^, *NKX2.8*^32^, *MMP9*^46^, and homologous recombination (HR) repair genes^47^. While most of the previous work on SMYD3 has focused on its implication in cancer, its role in normal differentiation programs has been largely ignored. The emerging interest in the impact of epigenetic regulators in differentiation has implicated members of the SMYD methyltransferase family in cardiac or skeletal myogenesis during development. These recent findings drove us to explore the molecular mechanisms and target gene pathways of SMYD3 during mammalian, skeletal muscle differentiation. We assessed the role of SMYD3 in mammalian myogenesis by analyzing the consequences of SMYD3 overexpression (SMYD3_OE_) and SMYD3 knockdown (SMYD3_KD_) on myoblast differentiation and gene expression. SMYD3 was originally linked to myogenesis by its requirement for cardiac and skeletal muscle development in zebrafish^35^. Our data provide the first evidence for a role of SMYD3 as a positive regulator of myogenic differentiation in mammals, as well as the first SMYD3-dependend transcriptomic analysis in differentiating myoblasts. We show that SMYD3_OE_ enhanced muscle differentiation in murine myoblasts, characterized by elevated expression of differentiation markers and increased myotubes size (Fig.1). Conversely, SMYD3_KD_ hampered the expression of differentiation markers and impaired myogenic differentiation *in vitro* (Fig. 2). Our transcriptome analysis identified SMYD3-dependent genes with important roles in skeletal muscle development, structure, organization and function (Fig.4). Furthermore, our loss-of-function analysis highlighted the conservation of SMYD3 function between human and mouse and supports an evolutionary conserved role for SMYD3 in myogenesis. Besides, we showed that the SMYD3 inhibitor BCI-121 significantly reduced C2C12 differentiation potential, supporting the importance of SMYD3 catalytic methyltransferase activity for proper myogenesis. Contrasting observations linking SMYD3 to muscle atrophy through the transcriptional activation of *c-Met* and *myostatin* genes^36^, suggest that SMYD3 could play diverse roles in normal development and in response to physiological stress, such as atrophy. Myogenesis requires a tightly-regulated balance between irreversible cell cycle exit, induction of muscle-specific genes, and functions of epigenetic regulators (for example, the SWI/SNF chromatin remodeling complex) ^40,48,49^. SMYD3’s role in cancer involves regulating cell proliferation and cell cycle progression^32,50,51^. So it was important to establish whether the muscle differentiation phenotypes could be explained by deregulation of cell cycle genes in SMYD3_OE_ or SMYD3_KD_ cells. Our proliferation analysis showed that SMYD3 is not primarily involved in the regulation of myoblasts cell cycle progression and cell cycle exit upon differentiation (Fig.3), thereby uncoupling myoblast proliferation from SMYD3-dependent differentiation roles.

Our study takes SMYD3 beyond cancer functions and links with recent reports implicating SMYD3 as a developmental regulator. For example, *Smyd3* expression is induced during human ES cell differentiation^52^ and SMYD3 contributes to early embryonic lineage commitment, where it participates in the blastocyst induction of pluripotency genes (*Oct4, Nanog, Sox2*), primitive endoderm markers (such as *Gata6*) and trophoectoderm markers (*Cdx2* and *Eomes*)^53^. Furthermore, SMYD3 maintains the self-renewal capacity of a subset of cancer stem cells (CSCs) by regulating the expression of the master stem cell regulator *ASCL2* ^54^. SMYD3 was detected in both undifferentiated and differentiated myoblasts and is not induced upon differentiation (Supplementary information Fig.S1). Furthermore, SMYD3 expression is detected in mesenchymal stem cells, from which the myogenic lineage derives (data not shown). Thus, it will be interesting to investigate if Smyd3 is required for early commitment to muscle cell identity and the de-repression of developmental regulators. Our transcriptomic analysis revealed that changes in SMYD3 levels primarily affect the expression of the key MRF myogenin and muscle-specific genes, suggesting that SMYD3 acts upstream of myogenin upon myoblast differentiation.

The activation of *myogenin* expression is required for the myogenic program, both during muscle development and post-natal muscle growth ^11,40,55^. *Myogenin* expression is directly regulated by MyoD, which can recognize E-box sequences in *myogenin* regulatory regions and can interact with chromatin remodeling factors and transcriptional regulators to drive expression ^6,56^. MyoD recruits histone acetylases (HATs) and the SWI/SNF remodeler complex to the *myogenin* locus ^18,19^. The KMT Set7/9 positively regulates myogenesis via interaction with MyoD and H3K4 methylation at regulatory regions of muscle genes ^22^. Myogenin can also regulate its own expression via a positive feed-back loop ^41^. Our transcriptome analysis showed that *myogenin* was the only MRF whose expression was significantly affected in both SMYD3_KD_ and SMYD3_OE_ cells, at 24h post-differentiation (Fig. 4). Furthermore, our ChIP experiments demonstrated that the deposition of the active transcription mark H3K4me3 at the *myogenin* locus is partially SMYD3-dependent, thus suggesting that SMYD3 is an additional epigenetic factor contributing to the transcriptional activation of *myogenin* upon myogenesis. However, it is possible that this regulation is through direct or indirect mechanisms. Indeed, we could not detect direct recruitment and binding of SMYD3 at the *myogenin* locus upon differentiation, nor evidence for a physical interaction with MyoD (unpublished data). In addition to its histones methylation activities, SMYD3 can facilitate mRNA transcription via other mechanisms, including direct interaction with Pol II and the transcriptional cofactors PC4 and BRD4^36,57^. Hence, we cannot exclude that the SMYD3 regulation of *myogenin* could also occur indirectly, via these alternative mechanisms.

Among the MRFs, myogenin is essential to guide terminal differentiation, myofiber formation and size^12^. Myoblasts fusion is guided by specific fusion proteins, such as the recently identified Myomaker and Myomerger ^58,59^, whose expression is under myogenin control. We observed substantially fewer multinucleated myotubes in SMYD3_KD_ cells compared to controls (Fig.2), whereas SMYD3_OE_ increased the number of MHC positive cells and the size of formed myotubes (Fig.1). Furthermore, our transcriptomic analysis detected SMYD3-dependent changes in the expression of genes involved in myoblast fusion, including the key fusion protein Myomaker. Thus, SMYD3 might regulate the expression of muscle-specific genes at the onset of differentiation, but also the later myoblast fusion processes necessary for correct terminal differentiation. Interestingly, *Myog*^*-/-*^ myoblasts also fail to correctly form myofibers, suggesting that impaired myoblast fusion in SMYD3_KD_ C2C12 might depend on the lack of myogenin^10,11^. Our transcriptome analysis highlighted that myogenin target genes are differentially expressed in SMYD3_KD_ and SMYD3_OE_ cells (Fig.4). It will be important to further investigate whether SMYD3 directly regulates the expression of genes involved in myoblast fusion, or whether this is an indirect effect of the deregulation of early myogenesis events, notably *myogenin* transcriptional activation.

Our data using myoblast models show that SMYD3 positively affects myogenesis. However, *Smyd3*^*-/-*^ mice are viable with no clear developmental defects^60^. This might be explained by compensation *in vivo* mediated by proteins with redundant functions. The SMYD protein family counts five different members which share a common conserved structure and some common histone targets, which may ensure redundancy during development ^61^. Furthermore, studies have implicated SMYD1-4 in cardiac and skeletal muscle development ^26,27^. SMYD1 is the only member showing a muscle-restricted pattern of expression^28,62^. We observed that *Smyd1* transcript levels were affected in both SMYD3_KD_ and SMYD3_OE_ cells (Supplementary information, Fig. S3). Previous studies revealed that *Smyd1* expression is directly regulated by myogenin^62^, suggesting that changes in Smyd1 expression are likely to be due to defective activation of *myogenin* upon SMYD3 expression modulation. Finally, we show that myogenin ectopic expression partially rescues terminal differentiation of SMYD3_KD_ C2C12, suggesting that SMYD3 and myogenin affect myoblasts fusion through overlapping and also distinct mechanisms. Our findings support a model in which SMYD3 can function upstream of a myogenin-dependent regulatory network, to regulate (directly or indirectly) endogenous *myogenin* expression. Both SMYD3_KD_ and SMYD3_OE_ phenotypes can be explained, at least partially, by the abnormal activation of endogenous myogenin expression, leading to inhibited or enhanced differentiation, respectively. However, only a subset of the deregulated genes in SMYD3_OE_ and SMYD3_KD_ cells can be explained by deregulated myogenin expression, highlighting the presence of a myogenin-independent SMYD3 regulatory pathway. In conclusion, our results highlight a new role for SMYD3 as a regulator of the myogenin pathway during muscle differentiation, potentially via its histone methyltransferase activity. We demonstrated that SMYD3 is a positive regulator of murine muscle differentiation, and this action is uncoupled from cell cycle exit, and we provided evidence for a conservation of its role in human myoblasts. Collectively, our data provide novel insights on the role of the lysine methyltransferase SMYD3 in the control of myogenic differentiation and muscle-specific gene expression.

## METHODS

### Cell lines and culture conditions

C2C12 mouse myoblasts were cultured in Growth medium (GM): Dulbecco’s modified Eagle’s medium (DMEM, Sigma) supplemented with 15% Fetal Calf Serum (FCS, Gibco) and 100 U/ml penicillin/streptomycin. Primary mouse myoblasts were cultured in plates coated with Gelatine 1%, in GM: DMEM supplemented with 20% FCS, 1% L-glutamine, 100 U/ml penicillin/streptomycin and 2% UltroserG. Muscle differentiation was induced by switching to serum-deprived differentiation medium (DM): DMEM, supplemented with 2% Horse Serum (Gibco), and 100 U/ml penicillin/streptomycin. Likewise, immortalized human myoblasts C25 were grown in GM: 1 Vol medium 199 (Gibco) + 4 Vol DMEM GlutaMAX (Gibco), 20% FCS, Fetuin: 25µg/ml, hEGF: 5ng/ml, bFGF: 0,5ng/ml, Insulin: 5 µg/ml, Dexamethasone: 0,2µg/ml, 100 U/ml penicillin/streptomycin. Differentiation was induced by switching to serum-deprived DM (DMEM + 100 U/ml penicillin/streptomycin + 10 μg/ml Insulin). All cell lines were cultured at 37°C and 5% CO_2_. Primary murine myoblasts were isolated from adult murine skeletal muscle and were kindly provided from Dr A. Pincini (Institut Cochin). C25 cells were provided by Dr A. Bigot (Institut de Myologie). All cell lines were cultured at 37°C and 5% CO2. The SMYD3 small-molecule inhibitor BCI-121 (Sigma, SML1817) was dissolved in DMSO at a concentration of 20 mg/ml. C2C12 cells were cultured for 24h in GM containing 100µM of BCI-121 or DMSO as a negative control. Cells when then switched to DM containing 100µM of BCI-121 or DMSO for 72 hrs. Protein samples were harvested at 0, 24, 48 and 72h post-differentiation upon BCI-121 treatment and analyzed by western blot for differentiation markers expression.

### Transfection and Gene Silencing

C2C12 cells, primary mouse myoblasts and human immortalized myoblasts were transfected with 60nM final of siRNAs (Sigma Aldrich) using LipoRNAiMAX (Invitrogen), according to the manufacturer’s instructions. Two consecutive transfections were performed, the first one on proliferative non confluent myoblasts and the second one 24h later, when myoblasts were induced to differentiation. The specific oligo sequences of siRNA are listed in Supplementary Table S6. Plasmids expressing MYOGENIN in pEMSV vector were kindly provided by the laboratory of Dr S. Ait-Si-Ali (CNRS, UMR7216, Paris) and transfected in C2C12 at 0.1 µg/cm^2^ using Lipofectamine2000 (Invitrogen) according to the manufacturer’s instructions.

### Construction of C2C12 cell line stably expressing HA-Flag-SMYD3

The cDNA encoding SMYD3 was PCR amplified and cloned into the pREV retroviral vector, kindly obtained from the laboratory of Dr S. Ait-Si-Ali (CNRS, UMR7216, Paris). The plasmid contains an epitope tag (3 HA- and 3 Flag-tag) in 5’ of the cloning site and a selection marker. C2C12 myoblasts stably expressing double-tagged Flag-HA-SMYD3 protein were established using retroviral transduction strategy as previously described ^48^. C2C12 cells stably integrating pREV-SMYD3 or pREV control vector were sub-cloned to obtain 100% positive clonal populations. Stable ectopic SMYD3 expression in these clones was validate by western blot and immunofluorescence analysis, using anti-HA and anti-FLAG antibodies. We selected two C2C12 positive clonal populations for our further experiments, called SMYD3 CL3 and SMYD3 CL5.

### CRISPRi design and Lentiviral Transduction

Human immortalized myoblasts C25 were transduced with lentiviral particles encoding CRISPRi system as previously described ^37^. Briefly, sgRNAs were designed to target near the TSS of the gene of interest (100bp upstream) and cloned into pLKO5-sgRNA-EFS-tRFP vector. pHR-SFFV-dCas9-BFP-KRAB plasmid was used for KRAB-dCas9 expression. To produce lentiviral particles, 293T cells were co-transfected with an envelope plasmid (pMD2.G-VSV-G), a packaging vector (psPAX2) and either the dCas9-KRAB or sgRNA expression vectors using calcium phosphate transfection. 48h after transfection, the medium containing viral particles was harvested and cleared of cell debris by filtering through a 0,45μm filter. Cleared supernatant was centrifuged at 25 000 rpm for 2h at 4°C. Concentrated virus was used to infect human myoblasts. Medium was changed after 24h. Cells were at first transfected with Krab-dCAS9 vector, and FACS sorted to obtain BFP positive cells. Stable cell lines integrating pHR-SFFV-dCas9-BFP-KRAB were then reinfected with sgRNAs. Efficient and stable SMYD3 silencing throughout cell passages and differentiation was confirmed at both RNA and protein levels. For sgRNA sequences, see Supplementary Table S6.

### RNA Extraction and RT-qPCR

Total RNA was extracted using NucleoSpin RNA Kit (Machery Nagel) or TRIzol (Invitrogen), following the manufacturer’s instructions. cDNA synthesis was performed with the Reverse Transcriptase Superscript III and IV (Invitrogen). Quantitative PCR amplification was performed with the ABI 7500 machine (Applied Biosystems) using the Sybr Green reagent (Applied Biosystems). See Supplementary TableS6 for primer sequences. The detection of a single product was verified by dissociation curve analysis. Relative quantities of mRNA were analysed using the Δ*C*_t_ method. *Gapdh* and *Rplp0* were used for normalization.

### Library preparation and sequencing for RNA-seq

Total RNA from two to three biological replicates was extracted using NucleoSpin RNA Kit (Machery Nagel) or TRIzol (Invitrogen) following the manufacturer’s instructions. RNA concentrations were obtained using nanodrop or a fluorometric Qubit RNA assay (Life Technologies, Grand Island, New York, USA). The quality of the RNA (RNA integrity number) was determined on the Agilent 2100 Bioanalyzer (Agilent Technologies, Palo Alto, CA, USA) as per the manufacturer’s instructions. To construct the libraries, 100 to 200ng of high-quality total RNA sample (RIN >8) was processed using TruSeq Stranded mRNA kit (Illumina) according to manufacturer instructions. Briefly, after purification of poly-A containing mRNA molecules, mRNA molecules are fragmented and reverse-transcribed using random primers. Replacement of dTTP by dUTP during the second strand synthesis will permit to achieve the strand specificity. Addition of a single A base to the cDNA is followed by ligation of Illumina adapters. Libraries were quantified by qPCR using the KAPA Library Quantification Kit for Illumina Libraries (KapaBiosystems, Wilmington, MA) and library profiles were assessed using the DNA High Sensitivity LabChip kit on an Agilent Bioanalyzer.

### RNA-seq analysis

Libraries were sequenced on an Illumina Nextseq 500 instrument using 75 base-lengths read V2 chemistry in a paired-end mode (between 17 and 80 million reads). Quality of FASTQ files was assessed using FASTQC software (version FastQC-v0.11.5). Reads were aligned using hisat2 ^63^ to the Mus_musculus.GRCm38.dna (version hisat2-2.1.0). Reads were then count using Featurecount (version subread-1.6.3) ^64^ and differential expression analysis was performed using DESeq2 package version 1.22.2 ^65^ using R software version 3.5.2. We corrected a batch effect (due to extraction method used) using DESeq2 with ∼ Condition + Extraction. We selected differentially expressed genes for which p value adjusted was < 0.01.

### Chromatin Immunoprecipitation (ChIP)

C2C12 cells were harvested with a cell scraper, washed twice in PBS, and fixed in PBS – 1% formaldehyde 10 minutes at room temperature. Formaldehyde was neutralized by adding glycine at 0.125 M final concentration for 5 minutes. Fixed cells were extensively washed in cold PBS. Chromatin was prepared by two subsequent extraction steps (10min, 4°C). First, cells were resuspended into cell lysis buffer (Hepes pH 7.8 25 mM, MgCl2 1.5 mM, KCl 10 mM, DTT 1 mM, NP-40 0.1%), incubated 10 min on ice followed by centrifugation (5 minutes, 2000 rpm). Nuclei were then resuspended in nuclear lysis buffer (Hepes pH 7.9 50 mM, NaCl 140 mM, EDTA 1 mM, Triton X100 1%, Na-deoxycholate 0.1%, SDS 0.5%), and sonicated on Bioruptor Power-up (Diagenode) to obtain genomic DNA fragments with a bulk size of 150–300 bp. Following centrifugation (10 min, 13000 rpm) the supernatant was used for immunoprecipitation. Immunoprecipitation was performed by incubating 2µg of H3K4me3 antibody (Diagenode pAb-030-050 - C15410030) or IgG (sc-2025) with 5µg of chromatin in IP buffer (Hepes pH 7.9 50 mM, NaCl 140 mM, EDTA 1 mM, Triton X100 1%) at 4°C overnight. The following day, protein-A/G beads (Thermo Fisher Scientific, 88802) were added to the antibodies-chromatin complexes and incubated at 4°C for 4 hours. Immunoprecipitates were washed three times with IP buffer, one time with wash buffer (Tris pH 8 20 mM, LiCl 250 mM, EDTA 1 mM, NP-40 0.5%, Na-deoxycholate 0.5%), and two times with elution buffer (Tris pH 8 20 mM, EDTA 1 mM). Then immunoprecipitated chromatin was eluted by incubating beads with extraction buffer supplemented with 1% SDS at 65°C. Chromatin was then reverse cross-linked by adding NaCl (200mM) and RNAseA (0.3µg/µL) and incubating at 65°C overnight. The proteins were digested by adding Proteinase K (0.2 µg/µL) and incubating 4 hours at 37°C. Finally, DNA was purified with DNA Extraction – Qiagen DNeasy kit. DNA was resuspended in water.

### Western Blot

Total proteins were extracted with Laemmli lysis buffer, sonicated (30 s ON/30 s OFF for 5 min) resolved on 10.5% acrylamide/bis-acrylamide SDS–PAGE gels and transferred to nitrocellulose membranes (Thermo Fisher Scientific) in transfer buffer. Protein transfer was assessed by Ponceau-red staining. Membranes were blocked in Tris-buffered saline pH 7.4 containing 0.1% Tween-20 and 5% milk for 1 h at room temperature. Incubations with primary antibodies were carried out at 4 °C overnight using the manufacturer recommended dilutions in Tris-buffered saline pH 7.4, 0.2% Tween-20. After 1 h incubation with an anti-rabbit or anti-mouse peroxidase-conjugated antibody (Jackson ImmunoResearch) at room temperature, proteins were detected by chemiluminescence using SuperSignal (Thermo Fisher Scientific), with Fusion v15.11 software or with the LI-COR Odyssey FC Imaging System and Image Studio Lite Software. The following antibodies were used: mouse anti-MyHC (R&D Systems, MAB4470), rabbit anti-histone H3 (Abcam, ab1791), rabbit anti-GAPDH (Sigma, G9545), monoclonal anti-Flag M2-Peroxidase (Sigma, A8592), rabbit anti-MCK (Kindly provided by Dr Slimane Ait-si-Ali, raised by Dr H Ito), rabbit anti-SMYD3 (Abcam, ab187149), rabbit anti-SMYD3 (Diagenode, C15410253-100), rabbit anti-Cyclin D1 (Abcam, ab 16663), rabbit anti-P21 (ab 109199), mouse anti-MYOGENIN (F5D ab 1835).

### Immunofluorescence and EdU staining

For each experiment, at the indicated time point, cells were washed with phosphate buffered saline (PBS) and fixed in PBS 3.7% formaldehyde for 15 min at room temperature. Cells were rinsed twice in PBS, permeabilized with PBS 0.5% Triton X-100 for 5 min and then blocked for 30 min with PBS 1% BSA to prevent non-specific staining. After three washes in PBS 0.2% Tween-20, cells were incubated with primary antibodies in 1% BSA at room temperature for 40 min −1h. After washing in PBS 0.2% Tween, cells were incubated for 30 min with secondary antibodies (1:500). Following washing in PBS 0.2% Tween, cells were stained with Hoechst or directly mounted using ProLong™ Gold Antifade Mountant with DAPI (Thermo Scientific). Images of immunofluorescence staining were photographed with a fluorescent microscope Leica Inverted 6000 or Zeiss, using Metamorph or ZEN software, respectively. As primary antibody we used anti-MyHC antibody (R&D Systems, MAB4470) and anti-MyoG antibody (F5D supernatant from DSHB). As secondary antibody Alexa Fluor 488 Goat anti-mouse or Cy3 Goat anti-mouse, of different IgG subtypes when co-staining for MyoG/MyHC. Fusion index was calculated as the percentage of nuclei contained in MyHC-positive myotubes, compared to the total number of nuclei. Myotubes diameter was measured along the myotube using ImageJ. Cell proliferation was assessed using EdU assay. C2C12 and stable clones were plated and induced to differentiation, or transfected with siRNAs before induction of differentiation. EdU incorporation was monitored in two culture conditions: GM or 24h in DM. EdU was added in cell culture media at a final concentration of 10µM during 30min (at 37°C). Cells were washed in PBS and fixed in PBS 3.7% formaldehyde for 15 min. After washing in PBS, cells were permeabilized in PBS Triton X-100 0.5% for 15min. Cells were washed three times in PBS Tween 0.2% and stained using the Click-iT® EdU Alexa Fluor® 488 kit (Thermo Scientific), following manufacturer’s instruction. Cells were incubated for 1h at room temperature, followed by three washes in PBS. Cells were then fixed in PBS 3.7% formaldehyde for 10 min, washed three times in PBS, permeabilized in PBS Triton X-100 0.5% for 5min. After washes in PBS, cells were stained with DAPI and analysed with a fluorescent microscope (Leica Inverted 6000). EdU and DAPI staining were counted in three independent experiment per condition. % EdU positive cells was calculated as the ration of EdU positive nuclei to DAPI-labeled nuclei.

### Statistical analysis

Statistical significance was assessed using the statistical package included in GraphPad Prism 7.0 software via Student’s t test or the analysis of variance (ANOVA) test (two-way ANOVA). Error bars show standard error means (SEM). P values of less than 0.05 were considered as statistically significant.

## Supporting information

Supplementary Informations

## Acknowledgements

We thank the GENOM’IC sequencing facility of the Institut Cochin for the preparation of RNA-Seq libraries and handling the sequencing procedure. We thank Xi Liu (Station Biologique de Roscoff) and Benjamin Saintpierre (GENOM’IC, Institut Cochin) for helping on the RNA-sequencing analysis. We thank the Institut the Myologie for providing C25 cell line and Dr A. Pincini (Institut Cochin) for primary mouse myoblasts. We acknowledge the ImagoSeine core facility of the Institut Jacques Monod (member of the France BioImaging, ANR-10-INBS-04), as well as the Epigenomics Core Facility and the Vectorology Platform of the UMR7216 ‘Epigenetics and Cell Fate’. We also thank Dr Slimane Ait-si-Ali (UMR7216, Paris, France) for the gift of the pREV-lentiviral vector. We thank colleagues at UMR7216 ‘Epigenetics and Cell Fate’ for helpful comments and suggestions during this study. This work was supported by the LabEx “Who Am I?” #ANR-11-LABX-0071 and the Université de Paris IdEx #ANR-18-IDEX-0001 funded by the French Government through its “Investments for the Future” program. Additional funding from AFMTELETHON 16146, and the Fondation ARC pour la Recherche sur le Cancer (ARC n°155029). JBW is a Senior member of the Institut Universitaire de France (IUF) and SM is a Junior member (2012ND 3369). RC is a recipient of a BioSPC PhD fellowship and Fondation pour la Recherche Médicale fellowship (FDT20170437242).

## Author Contributions Statement

SM and JBW conceived the study and sought funding.

RC, MP, AD, EF and SM performed the experiments.

AS and AB provided reagents and experimental expertise.

RC, JBW and SM analyzed results, developed the scientific conclusions and wrote the manuscript.

## Additional Information

### Supplementary information

accompanies this paper.

## Data availability

All data generated or analyzed during this study are included in this published article. The RNA-Sequencing data have been deposited to the GEO with the dataset identifier GSE137954.

