## Supplementary Informations for "The SMYD3 methyltransferase promotes myogenesis by activating the myogenin regulatory network"

Figure S1

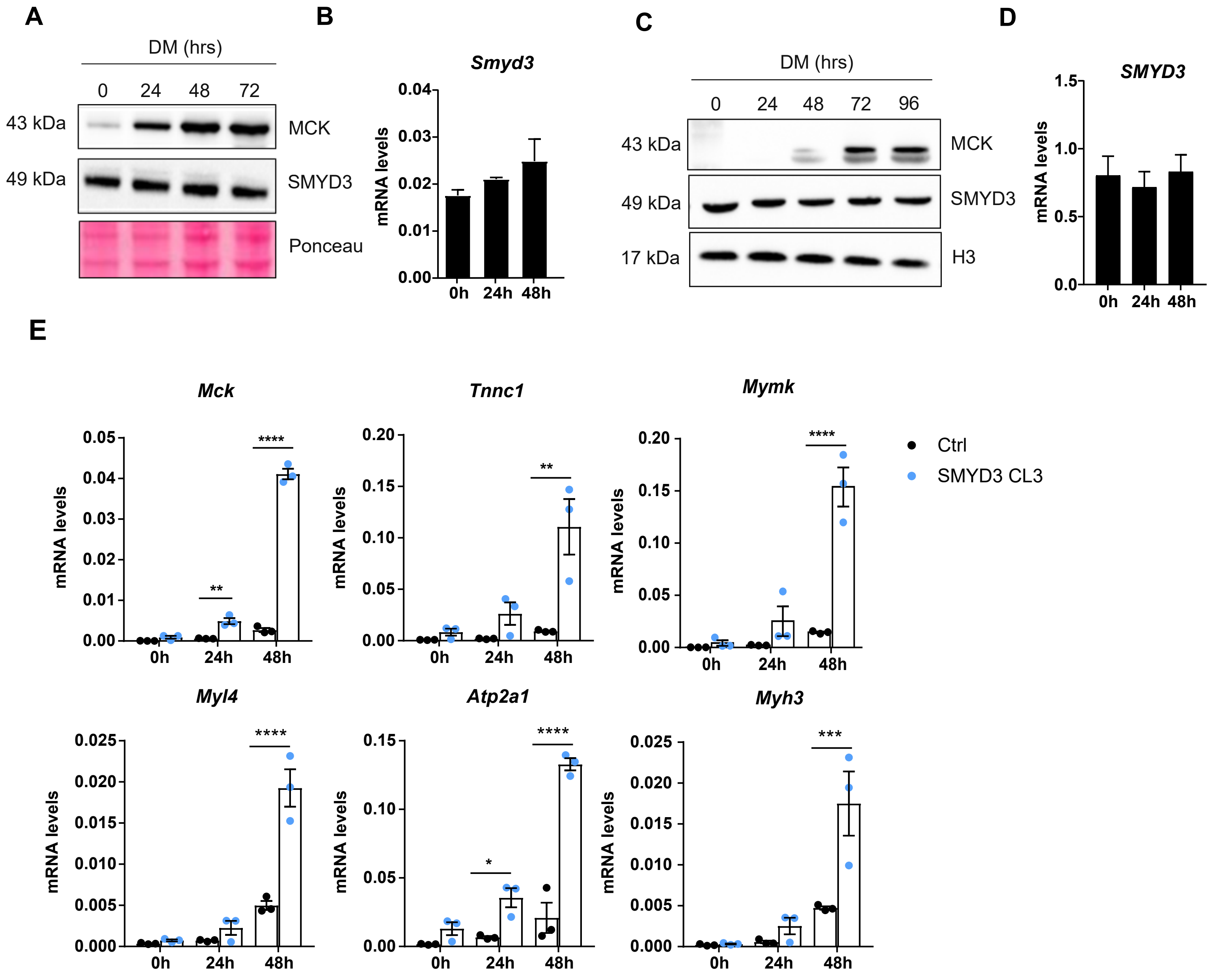
A
DM (hrs)
0
24
48
72
43 kDa
MCK
49 kDa
SMYD3
Ponceau
B
Smyd3
mRNA levels
0h
24h
48h
C
DM (hrs)
0
24
48
72
96
43 kDa
MCK
49 kDa
SMYD3
17 kDa
H3
D
SMYD3
mRNA levels
E
Mck
Tnnc1
Mymk
Myl4
Atp2a1
Myh3
Ctrl
SMYD3 CL3

#### Figure S2

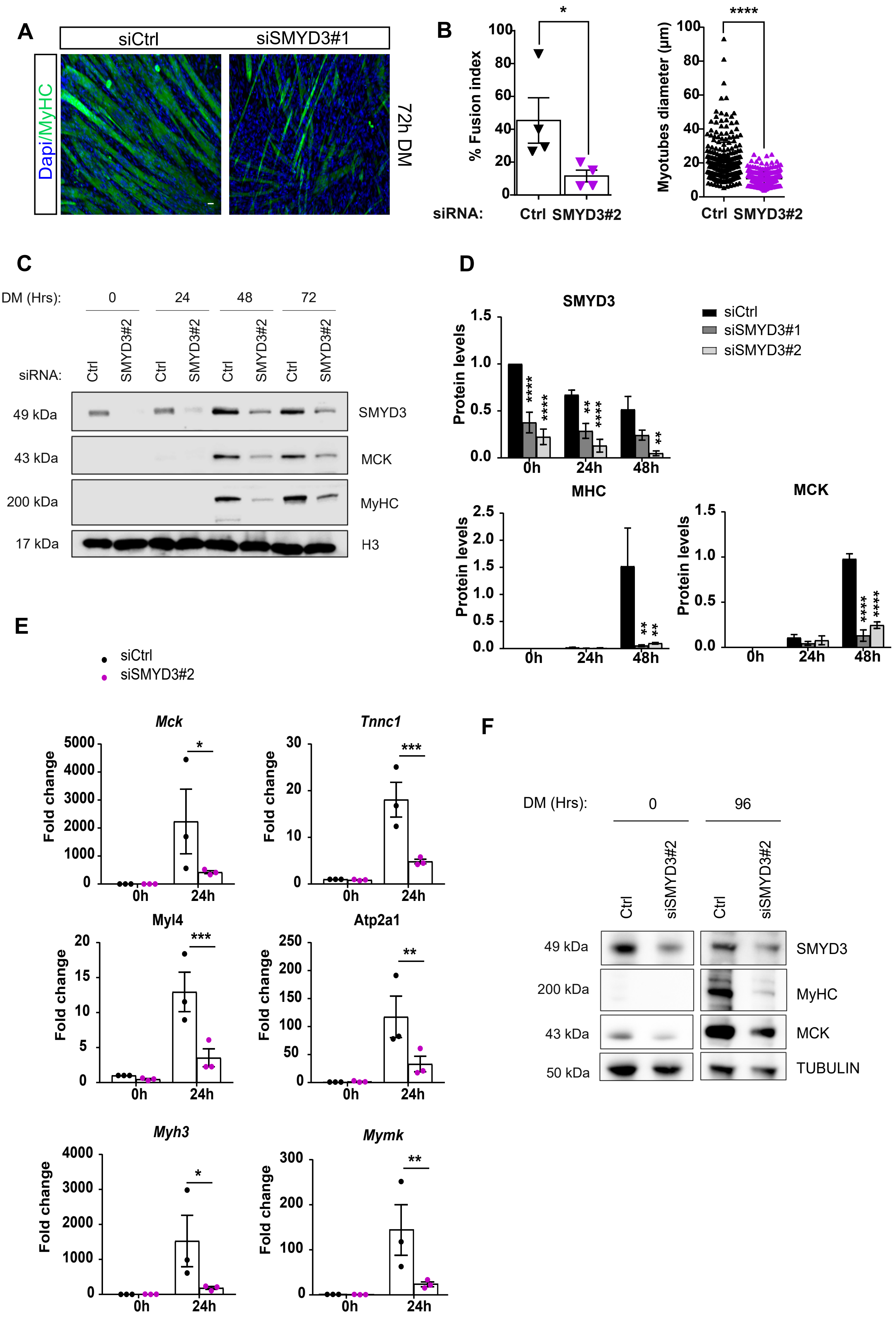

A
siCtrl
siSMYD3#1
Dapi/MyHC
72h DM
B
% Fusion index
siRNA:
Ctrl
SMYD3#2
*
Myotubes diameter (µm)
****
C
DM (Hrs):
0
24
48
72
siRNA:
Ctrl
SMYD3#2
49 kDa
43 kDa
200 kDa
17 kDa
SMYD3
MCK
MyHC
H3
D
SMYD3
MHC
MCK
Protein levels
siCtrl
siSMYD3#1
siSMYD3#2
0h
24h
48h
E
siCtrl
siSMYD3#2
Mck
Tnnc1
Myl4
Atp2a1
Myh3
Mymk
Fold change
F
DM (Hrs):
0
96
Ctrl
siSMYD3#2
49 kDa
200 kDa
43 kDa
50 kDa
SMYD3
MyHC
MCK
TUBULIN

### Figure S3

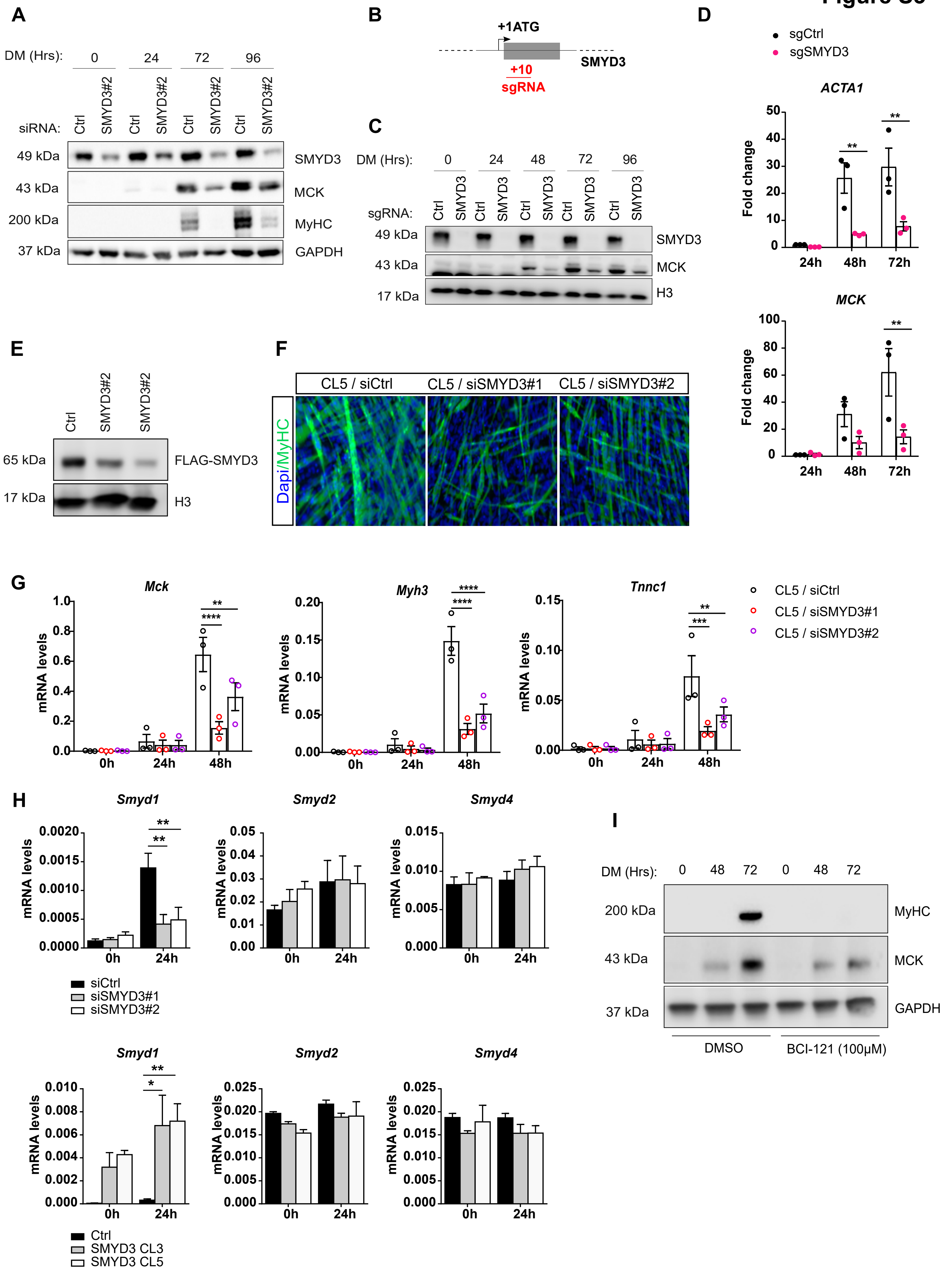

A
DM (Hrs): 0 24 72 96
siRNA: Ctrl SMYD3#2 Ctrl SMYD3#2 Ctrl SMYD3#2 Ctrl SMYD3#2
49 kDa SMYD3
43 kDa MCK
200 kDa MyHC
37 kDa GAPDH
B
+1ATG
+10
sgRNA
SMYD3
C
DM (Hrs): 0 24 48 72 96
sgRNA: Ctrl SMYD3 Ctrl SMYD3 Ctrl SMYD3 Ctrl SMYD3 Ctrl SMYD3
49 kDa SMYD3
43 kDa MCK
17 kDa H3
D
sgCtrl
sgSMYD3
ACTA1
MCK
Fold change
24h 48h 72h
E
Ctrl SMYD3#2 SMYD3#2
65 kDa FLAG-SMYD3
17 kDa H3
F
CL5 / siCtrl
CL5 / siSMYD3#1
CL5 / siSMYD3#2
Dapi/MyHC
G
Mck
Myh3
Tnnc1
mRNA levels
0h 24h 48h
CL5 / siCtrl
CL5 / siSMYD3#1
CL5 / siSMYD3#2
H
Smyd1
Smyd2
Smyd4
mRNA levels
0h 24h
siCtrl
siSMYD3#1
siSMYD3#2
Ctrl
SMYD3 CL3
SMYD3 CL5
I
DM (Hrs): 0 48 72 0 48 72
200 kDa MyHC
43 kDa MCK
37 kDa GAPDH
DMSO
BCI-121 (100μM)

### Figure S4

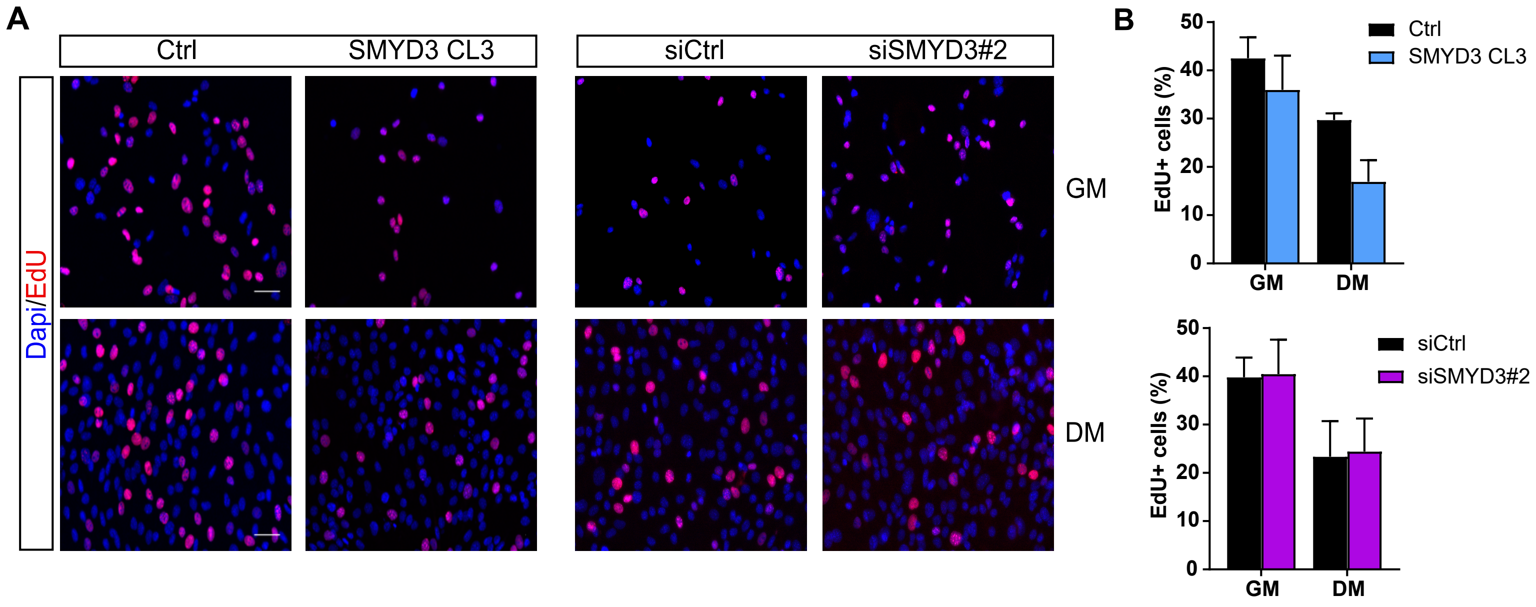
A
Ctrl
SMYD3 CL3
siCtrl
siSMYD3#2
Dapi/EdU
GM
DM
B
EdU+ cells (%)
50
40
30
20
10
0
GM
DM
Ctrl
SMYD3 CL3
siCtrl
siSMYD3#2

Figure S5

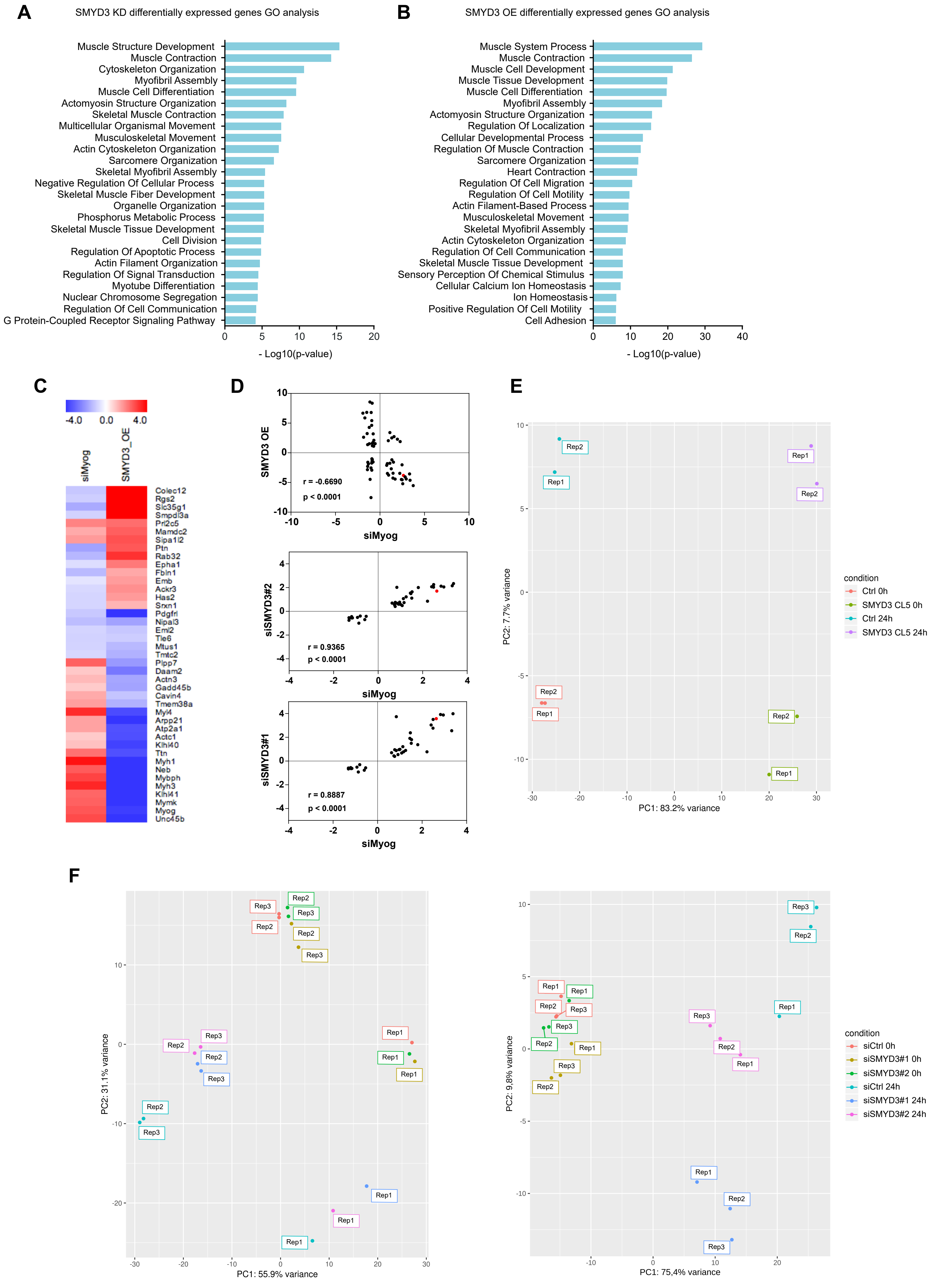
A
SMYD3 KD differentially expressed genes GO analysis
Muscle Structure Development
Muscle Contraction
Cytoskeleton Organization
Myofibril Assembly
Muscle Cell Differentiation
Actomyosin Structure Organization
Skeletal Muscle Contraction
Multicellular Organismal Movement
Musculoskeletal Movement
Actin Cytoskeleton Organization
Sarcomere Organization
Skeletal Myofibril Assembly
Negative Regulation Of Cellular Process
Skeletal Muscle Fiber Development
Organelle Organization
Phosphorus Metabolic Process
Skeletal Muscle Tissue Development
Cell Division
Regulation Of Apoptotic Process
Actin Filament Organization
Regulation Of Signal Transduction
Myotube Differentiation
Nuclear Chromosome Segregation
Regulation Of Cell Communication
G Protein-Coupled Receptor Signaling Pathway
0 5 10 15 20
- Log10(p-value)
B
SMYD3 OE differentially expressed genes GO analysis
Muscle System Process
Muscle Contraction
Muscle Cell Development
Muscle Tissue Development
Muscle Cell Differentiation
Myofibril Assembly
Actomyosin Structure Organization
Regulation Of Localization
Cellular Developmental Process
Regulation Of Muscle Contraction
Sarcomere Organization
Heart Contraction
Regulation Of Cell Migration
Regulation Of Cell Motility
Actin Filament-Based Process
Musculoskeletal Movement
Skeletal Myofibril Assembly
Actin Cytoskeleton Organization
Regulation Of Cell Communication
Skeletal Muscle Tissue Development
Sensory Perception Of Chemical Stimulus
Cellular Calcium Ion Homeostasis
Ion Homeostasis
Positive Regulation Of Cell Motility
Cell Adhesion
0 10 20 30 40
- Log10(p-value)
C
-4.0 0.0 4.0
siMyog
SMYD3_OE
Colec12
Rgs2
Slc35g1
Smpdl3a
Prl2c5
Mamdc2
Sipa1l2
Ptn
Rab32
Epha1
Fbln1
Emb
Ackr3
Has2
Srxn1
Pdgfrl
Nipal3
Eml2
Tle6
Mtus1
Tmtc2
Plpp7
Daam2
Actn3
Gadd45b
Cavin4
Tmem38a
Myl4
Arpp21
Atp2a1
Actc1
Klhl40
Ttn
Myh1
Neb
Mybph
Myh3
Klhl41
Mymk
Myog
Unc45b
D
SMYD3 OE
siMyog
r = -0.6690
p < 0.0001
siSMYD3#2
siMyog
r = 0.9365
p < 0.0001
siSMYD3#1
siMyog
r = 0.8887
p < 0.0001
E
PC2: 7.7% variance
PC1: 83.2% variance
condition
Ctrl 0h
SMYD3 CL5 0h
Ctrl 24h
SMYD3 CL5 24h
F
PC2: 31.1% variance
PC1: 55.9% variance
PC2: 9.8% variance
PC1: 75,4% variance
condition
siCtrl 0h
siSMYD3#1 0h
siSMYD3#2 0h
siCtrl 24h
siSMYD3#1 24h
siSMYD3#2 24h

### Figure S6

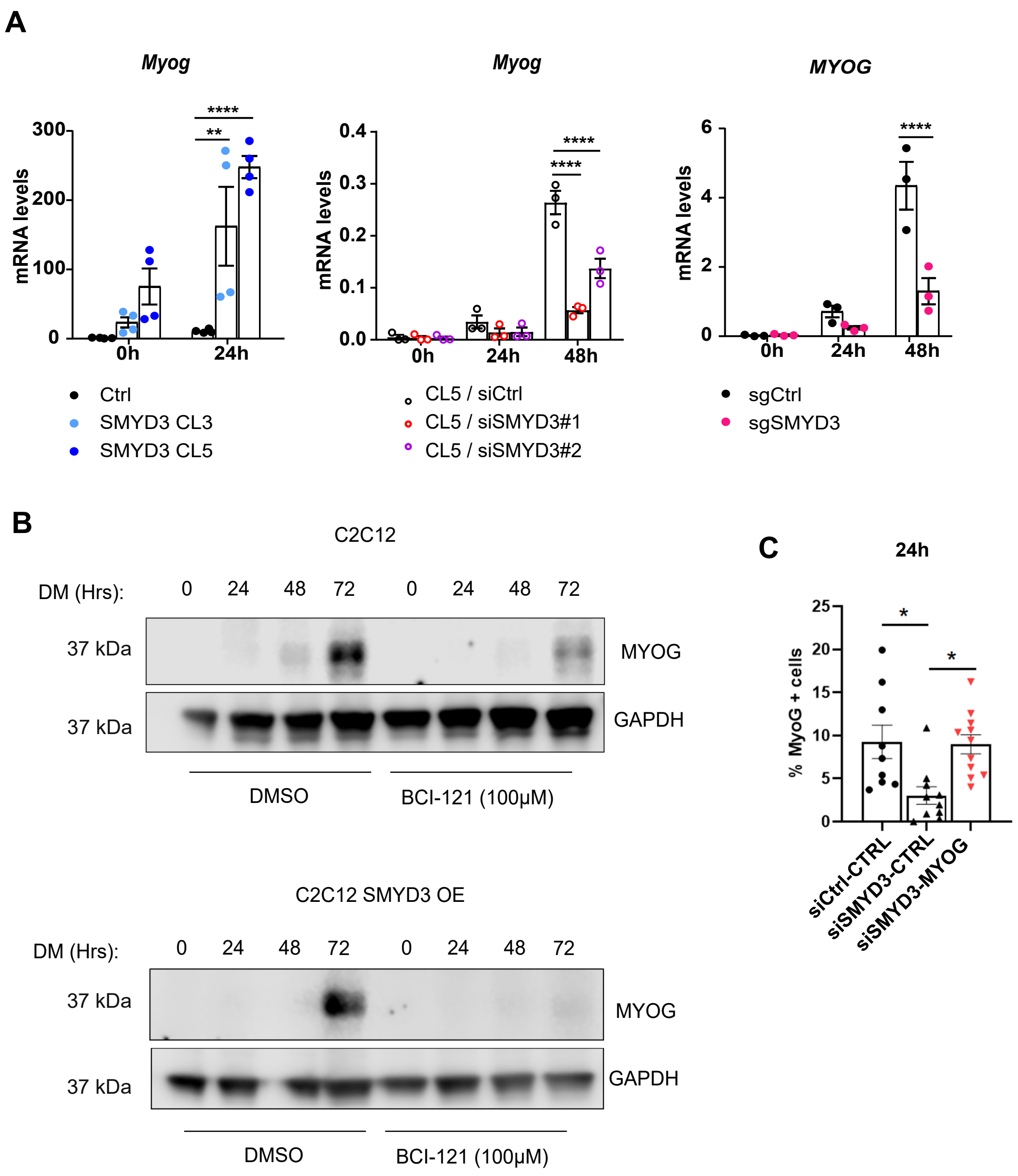

A
Myog
mRNA levels
300
200
100
0
0h
24h
****
**
Ctrl
SMYD3 CL3
SMYD3 CL5
Myog
mRNA levels
0.4
0.3
0.2
0.1
0.0
0h
24h
48h
****
****
CL5 / siCtrl
CL5 / siSMYD3#1
CL5 / siSMYD3#2
MYOG
mRNA levels
6
4
2
0
0h
24h
48h
****
sgCtrl
sgSMYD3
B
C2C12
DM (Hrs):
0 24 48 72 0 24 48 72
37 kDa
MYOG
37 kDa
GAPDH
DMSO
BCI-121 (100μM)
C2C12 SMYD3 OE
DM (Hrs):
0 24 48 72 0 24 48 72
37 kDa
MYOG
37 kDa
GAPDH
DMSO
BCI-121 (100μM)
C
24h
% MyoG + cells
25
20
15
10
5
0
*
*
siCtrl-CTRL
siSMYD3-CTRL
siSMYD3-MYOG

### Figure S7

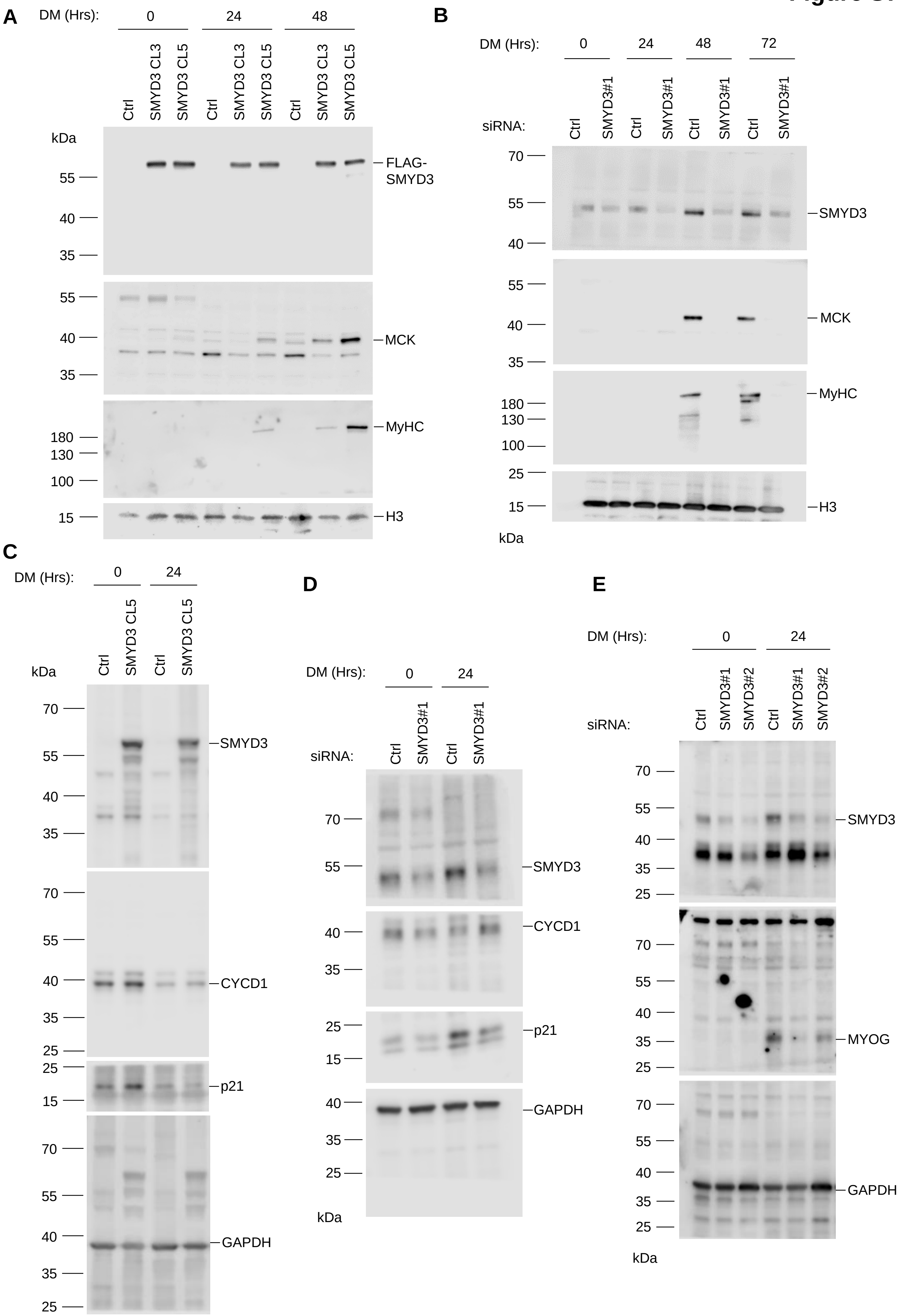

A
DM (Hrs):
0
24
48
Ctrl
SMYD3 CL3
SMYD3 CL5
kDa
55
40
35
FLAG-
SMYD3
MCK
180
130
100
MyHC
15
H3
B
DM (Hrs):
0
24
48
72
siRNA:
Ctrl
SMYD3#1
70
SMYD3
MCK
MyHC
25
C
CYCD1
p21
GAPDH
D
E
SMYD3#2
MYOG

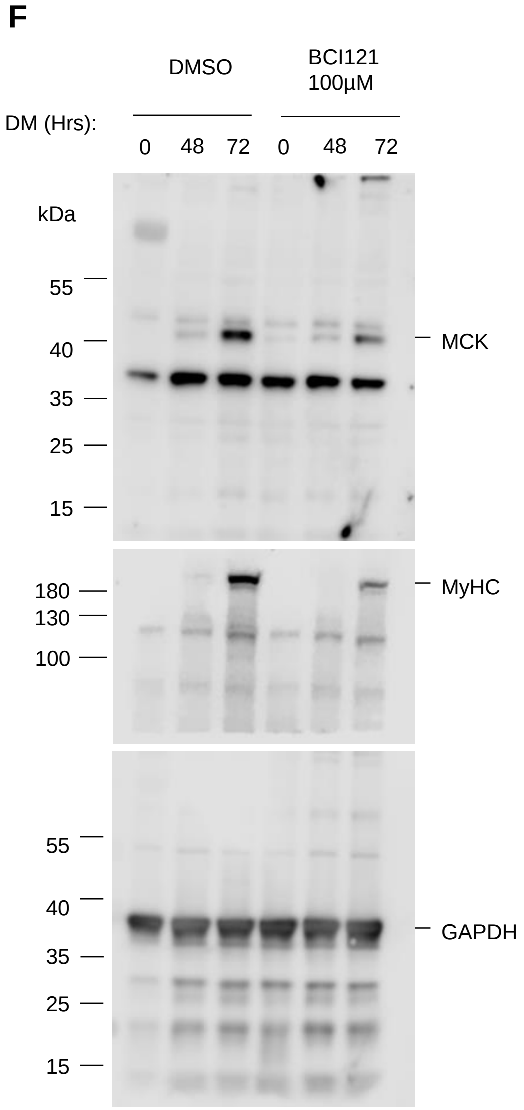
F
DMSO
BCI121
100µM
DM (Hrs):
0
48
72
0
48
72
kDa
55
40
35
25
15
MCK
180
130
100
MyHC
55
40
35
25
15
GAPDH

**Figure S1. SMYD3 is expressed in undifferentiated and differentiated myoblasts and its overexpression enhances myogenesis**

**A)** Time course of SMYD3 protein expression during terminal differentiation of murine myoblasts. Muscle Creatine Kinase (MCK) is used as differentiation marker, Ponceau red as loading control.

**B)** qPCR analysis of *Smyd3* expression during terminal differentiation of murine myoblasts. mRNA levels were normalized to *Gapdh* Ct values at the indicated time points. Data shown as means ± SEM of three independent experiments.

**C)** Time course of SMYD3 protein expression during terminal differentiation of human immortalized myoblasts. MCK is used as differentiation marker. H3 is a loading control.

**D)** qPCR analysis of *SMYD3* expression during terminal differentiation of human immortalized myoblasts. mRNA levels were normalized to *GAPDH* Ct values at the indicated time points. Data shown as means ± SEM of three independent experiments.

**E)** qPCR analysis on Ctrl and SMYD3 CL3 $SMYD3_{OE}$ cells at 0h, 24h and 48h in DM. mRNA levels of specific differentiation markers (*Mck, Tnnc1, Myl4, Myh3, Atp2a1, Mymk*) were normalized to *Gapdh* and *Rplp0* Ct values at the indicated timepoints. Graphs present means ± SEM of four independent experiments. ANOVA, * $p < 0.05$, ** $p < 0.01$, *** $p< 0.001$, **** $p< 0.0001$ vs. control respectively.

**Figure S2. SMYD3 silencing delays skeletal myogenesis**

**A)** MyHC immunofluorescence analysis performed in differentiated in C2C12 myoblasts after transfection with control or *Smyd3* siRNAs. Cells were transferred to differentiation medium and stained with MyHC antibody at the indicated timepoints. Cells were Hoechst stained prior to immunofluorescence analysis. Scale bar, 50µm.

**B)** Left panel: quantification of fusion index of four independent experiments calculated as percentage of nuclei within MyHC-expressing myotubes at 72h in presence of the various siRNA. Data shown as mean ± SEM. Mann-Whitney, * $p < 0.05$. Right panel: quantification of myotubes diameter of >100 myotubes in four independent experiments, calculated within MyHC-expressing myotubes. Data are presented as average ± SEM. Mann-Whitney, **** $p < 0.0001$.

**C)** Time course of protein expression during terminal differentiation of $SMYD3_{KD}$ cells. Cellular extracts were analyzed by western blot at 0h, 24h, 48h and 72h in DM with antibodies against SMYD3, muscle creatine kinase (MCK), and myosin heavy chain (MyHC). H3 is a loading control.

**D)** Quantification of SMYD3, MCK and MyHC protein levels from 2 to 4 independent western blot. Data shown as mean ± SEM. Significative p-values (ANOVA) are indicated. ** $p < 0.01$ **** $p<0.001$.

**E)** qPCR analysis on C2C12 cells after transfection with control or *Smyd3* siRNAs. mRNA levels of specific differentiation markers (*Mck, Tnnc1, Myl4, Myh3, Atp2a1, Mymk*) were normalized to *Gapdh* and *Rplp0* Ct values at the indicated time points. Data shown as mean fold change ± SEM of three independent experiments. ANOVA, * $p < 0.05$, ** $p < 0.01$ **** $p<0.001$ vs. control respectively.

**F)** Reduced protein levels of MyHC and MCK during terminal differentiation of primary mouse myoblasts after transfection with control or *Smyd3* siRNAs. Cellular extracts were analyzed by western blot at 0h and 96h in DM with antibodies against SMYD3, Muscle Creatine Kinase (MCK), and myosin heavy chain (MyHC). Tubulin is a loading control.

**Figure S3. SMYD3 knockdown delays skeletal myogenesis in human myoblasts and in SMYD3$_{OE}$ cells**

**A)** Time course of protein expression during terminal differentiation of human immortalized myoblasts in which control or *SMYD3* siRNAs were delivered in GM. Cellular extracts were analyzed by western blot at the indicated time points in DM with antibodies against SMYD3, Muscle Creatine Kinase (MCK), and myosin heavy chain (MyHC). GAPDH is a loading control.

**B)** Schematic of 5' genomic region of *SMYD3* gene: coding sequence (grey box), start codon (arrow, ATG), position and size of sgRNA used to silence *SMYD3* expression using CRISPRi.

**C)** Representative time course of protein expression during terminal differentiation of human myoblasts cell lines expressing sgCtrl and sgSMYD3. Myoblasts were cultured in growth medium (GM) until they reached confluence, and then induced to differentiate by switching to differentiation medium (DM). Cellular extracts were analyzed by western blot at the indicated time points with antibodies against SMYD3 and Muscle Creatine Kinase (MCK). H3 is a loading control.

**D)** qPCR analysis of mRNA levels of the differentiation markers *ACTA1* and *MCK* in sgCtrl and sgSMYD3 human myoblasts cell lines at 24h, 48h and 72h in DM. mRNA levels were normalized to *GAPDH* Ct values. Data shown as mean fold change ± SEM of three independent experiments. ANOVA, ** $p < 0.01$ vs. control.

**E)** Western blot analysis of SMYD3-FLAG protein levels in SMYD3 CL5, after transfection with control or *Smyd3* siRNAs. H3 is used as loading control.

**F)** Immunofluorescence analysis of MyHC staining after SMYD3 silencing in SMYD3-CL5 cells.

**G)** qPCR analysis of mRNA levels of the differentiation markers *Mck*, *Myh3* and *Tnnc1* in SMYD3-CL5 cells after SMYD3 silencing. mRNA levels were normalized to *Gapdh* Ct values. Data shown as mean fold change ± SEM of three independent experiments. ANOVA, ** $p < 0.01$ **** $p< 0.001$ vs. control respectively.

**H)** qPCR analysis of mRNA levels of *Smyd1*, *Smyd2* and *Smyd4* after SMYD3 knockdown (upper panel) or stable overexpression (lower panel). mRNA levels were normalized to *Gapdh* and *Rplp0* Ct values. Data shown as mean fold change ± SEM of three independent experiments. ANOVA, * $p<0.05$, ** $p < 0.01$ vs. control respectively.

**I)** SMYD3$_{OE}$ cells (SMYD3 CL5) were treated with control DMSO or 100µM of BCI-121 and differentiated for 72h. Cellular extracts were analyzed by western blot at the indicated time points with antibodies against muscle creatine kinase (MCK), and myosin heavy chain (MyHC). GAPDH is a loading control.

**Figure S4 SMYD3 is not involved in the regulation of myoblasts proliferation and cell cycle**

**A)** EdU staining was performed in $SMYD3_{OE}$ cells (SMYD3 CL3) and control cells (Ctrl) (left panel) and in $SMYD3_{KD}$ (SMYD3#2) or a scramble sequence (siCtrl) (right panel). In both cases, cells were labeled with EdU either in GM or after 24h in DM. Pictures are one representative of 3 independent experiments. EdU (red) and DAPI (blue) staining (scale bar: 50μm).

**B)** Percentage of EdU positive nuclei/DAPI labeled nuclei in randomly selected fields, in $SMYD3_{OE}$ or $SMYD3_{KD}$ cells, compared with relative controls. Error bars represent SEM from quantifications of 3 independent experiments.

**Figure S5. RNA-seq differential expression analysis of $SMYD3_{KD}$ and $SMYD3_{OE}$ C2C12**

**A and B)** PANTHER GO enrichment analysis tool was used to identify significant GO terms (Biological Process) enriched in the list of differentially expressed genes in $SMYD3_{KD}$ cells and $SMYD3_{OE}$ cells at 24h post-differentiation.

**C)** Heat map showing the expression profiles of common DE transcripts in siMyog (from the study of Liu et al., 2010) and $SMYD3_{OE}$ cells, at 24h of differentiation.

**D)** Scatterplot graphs and correlation analysis for common DE transcripts in siMyog microarray (Liu et al, 2010) and $SMYD3_{OE}$ and $SMYD3_{KD}$ RNA-seq in C2C12 cells at 24h post-differentiation (expression levels for the two siRNAs used in this study are represented separately). On the axes, the log2(fold-change) values (adjusted p-value > 0.01). Each point is a gene, *Myogenin* is indicated in red on each plot. Pearson correlation coefficient (r) and p-value for each comparison are indicated on the graph.

**E)** Principal component analysis (PCA) results for $SMYD3_{OE}$ experiments, each point represents a different replicate.

**F)** Principal component analysis (PCA) results for $SMYD3_{KD}$ experiments, each point represents a different replicate. Left panel: we could observe a batch effect due to RNA extraction method between Replicate 1 and Replicates 2/3 that was corrected as described in Material and Methods. Right panel: PCA results after correction.

**Figure S6. SMYD3 regulates *myogenin* transcriptional activation**

**A)** Relative expression levels of *myogenin* transcripts were monitored by qPCR in differentiating $SMYD3_{OE}$ cells (SMYD3 CL3 and CL5) and control cells (Ctrl) or in $SMYD3_{OE}$ cells after transfection with control or *Smyd3* siRNAs, and in sgCtrl and sgSMYD3 human myoblasts cell lines. Transcript levels were analyzed at the indicated time points for each condition. mRNA levels were normalized to *Gapdh* and *Rplp0* Ct values. Data shown as means ± SEM of three independent experiments. ANOVA, ** $p < 0.01$ **** $p < 0.0001$ vs. control respectively.

**B)** C2C12 cells (top) and $SMYD3_{OE}$ cells (SMYD3 CL5) (bottom) were treated with control DMSO or 100µM of BCI-121 and differentiated for 72h. Myogenin expression was analyzed by western blot at the indicated time points in DM. GAPDH is a loading control.

**C)** The percentage of myogenin positive cells was calculated by immunofluorescence staining as the percentage of positive nuclei / the total number of nuclei per field, in siRNA-treated C2C12 cells at 24h in DM, following overexpression of myogenin or a control vector. Data shown as means ± SEM. ANOVA, ** $p < 0.01$, * $p < 0.05$.

**Figure S7. Original blots**

**A)** Original blot of Fig.1C. Membranes were cut before incubation with the antibody.

**B)** Original blot of Fig.2C. Membranes were cut before incubation with the antibody.

**C)** Original blot of Fig.3C. Membranes were cut before incubation with the antibody.

**D)** Original blot of Fig.3D. Membranes were cut before incubation with the antibody.

**E)** Original blot of Fig.5B. Membranes were cut before incubation with the antibody.

**F)** Original blot of Fig.2E Membranes were cut before incubation with the antibody.

#### SUPPLEMENTARY TABLES

**Table S1. List of differentiation-dependent genes deregulated in $SMYD3_{KD}$ C2C12 myoblasts at 24h post-differentiation**

This table includes log2 fold change (log2FC) and p-value adjusted (padj) of the differentially expressed genes in both siRNAs used in this study (siSMYD3#1 and siSMYD3#2), at 24h post-differentiation compared to the control (Ctrl).

**Table S2. List of differentiation-dependent genes deregulated in $SMYD3_{OE}$ C2C12 myoblasts at 24h post-differentiation**

This table includes log2 fold change (log2FC) and p-value adjusted (padj) of the differentially expressed genes in overexpressing cells (SMYD3 CL5), at 24h post-differentiation compared to the control (Ctrl).

**Table S3. List of common deregulated genes between $SMYD3_{KD}$ and $SMYD3_{OE}$ C2C12 myoblasts at 24h post-differentiation**

This table includes log2 fold change (log2FC) and p-value adjusted (padj) of the differentially expressed genes in both siRNAs used in this study (siSMYD3#1 and siSMYD3#2) and in overexpressing cells (SMYD3 CL5), at 24h post-differentiation compared to the control (Ctrl).

**Table S4. Commonly deregulated genes between $SMYD3_{KD}$, $SMYD3_{OE}$ and siMyog C2C12 myoblasts at 24h post-differentiation**

The Microsoft Excel workbook contains two tabs. The first one (siCtrl_siMyog_24h) contains expression profiles of C2C12 myoblasts after siRNA-mediated knockdown of *myogenin*, compared with a control siRNA, at 24h post-differentiation. These data were obtained from re-analysis of raw data generated by Liu et al., 2010 available on the Gene Expression Omnibus GEO (#GSE19967). The second tab (Common_SMYD3KD) contains expression profiles of commonly deregulated genes between siMyog and siSMYD3 C2C12 myoblasts, at 24h post-differentiation (both siRNAs used in this study are reported). The third tab (Common_SMYD3OE) contains expression profiles of commonly deregulated genes between siMyog and SMYD3-overepxressing C2C12 myoblasts, at 24h post-differentiation.

**Table S5. Number of reads detected in RNA-seq experiment per sample**

**Table S6. Sequence of all DNA and RNA oligonucleotides used in this work**
